# The C-terminal tail of the bacterial translocation ATPase SecA modulates its activity

**DOI:** 10.1101/389460

**Authors:** Mohammed Jamshad, Timothy J. Knowles, Scott A. White, Douglas G. Ward, Fiyaz Mohammed, Kazi Rahman, Gareth W. Hughes, Günter Kramer, Bernd Bukau, Damon Huber

**Affiliations:** Institute for Microbiology and Infection, School of Biosciences, University of Birmingham, Edgbaston, Birmingham, UK; Institute of Cancer and Genomic Sciences, University of Birmingham, Edgbaston, Birmingham, UK; Institute of Immunology and Immunotherapy, University of Birmingham, Edgbaston, Birmingham, UK; Center for Molecular Biology of Heidelberg University (ZMBH); German Cancer Research Center (DKFZ), ZMBH-DKFZ Alliance, Heidelberg, Germany

## Abstract

In bacteria, the translocation of a subset of proteins across the cytoplasmic membrane by the Sec machinery requires SecA. Although SecA can recognise nascent polypeptides, the mechanism of cotranslational substrate protein recognition is not known. Here, we investigated the role of the C-terminal tail (CTT) of SecA, which consists of a flexible linker (FLD) and a small metal-binding domain (MBD), in its interaction with nascent polypeptides. Phylogenetic analysis and ribosome binding experiments indicated that the MBD interacts with 70S ribosomes. Disruption of the entire CTT or the MBD alone had opposing effects on ribosome binding, substrate-protein binding, ATPase activity and in vivo function. Autophotocrosslinking, mass spectrometry, x-ray crystallography and small-angle x-ray scattering experiments provided insight into the CTT-mediated conformational changes in SecA. Finally, photocrosslinking experiments indicated that binding of SecA to substrate protein affected its interaction with the ribosome. Taken together, our results suggest a mechanism for substrate protein recognition.

**Impact Statement:** SecA is an evolutionarily conserved ATPase that is required for the translocation of a subset of proteins across the cytoplasmic membrane in bacteria. We investigated how SecA recognises its substrate proteins at the ribosome as they are still being synthesised (i.e. cotranslationally).

## Introduction

In *Escherichia coli*, approximately a quarter of all newly synthesised proteins are transported across the cytoplasmic membrane by the Sec machinery (Cranford Smith & Huber, 2018; Van den Berg et al., 2004). Of these, the majority (∼65%) require the assistance of SecA, a evolutionarily conserved and essential motor ATPase that drives protein translocation in bacteria (Cranford Smith & Huber, 2018). The catalytic core of SecA (amino acids ∼1-832 in *E. coli*) contains five domains (**supplemental figure S1**): nucleotide binding domain-1 (NBD1; amino acids 9-220 & 378-411), nucleotide binding domain-2 (NBD2; 412-620), the polypeptide crosslinking domain (PPXD; 221-377), the α-helical scaffold domain (HSD; 621-672 & 756-832) and the α-helical wing domain (HWD; 673-755). Binding and hydrolysis of ATP at the interface of NBD1 and NBD2 cause conformational changes in the HSD and HWD, which drive translocation (Collinson, Corey, & Allen, 2015; Cranford Smith & Huber, 2018). In addition, the PPXD undergoes a large conformational change, swinging from a position near the HWD (the “closed” conformation) to a position near NBD2 (the “open” conformation) (Chen, Bauer, Rapoport, & Gumbart, 2015; Zimmer, Nam, & Rapoport, 2008; Zimmer & Rapoport, 2009). SecA binds to substrate protein in the groove formed between the PPXD and the two NBDs, and the PPXD serves as a “clamp” that locks unfolded substrate proteins into this groove when it is in the open conformation (Zimmer & Rapoport, 2009).

SecA can recognise proteins that are destined for translocation across the cytoplasmic membrane while they are still being synthesised (i.e. cotranslationally) (Huber et al., 2017; Huber et al., 2011). Previous work indicates that SecA binds to the ribosome and that ribosome binding facilitates its interaction with nascent chains (Chun & Randall, 1994; Huber et al., 2017; Huber et al., 2011). The binding site for SecA on the ribosome includes ribosomal protein uL23, which is located adjacent to the opening of the polypeptide exit channel (Huber et al., 2011). Binding is mediated by the N-terminal α-helix of SecA and the N-terminal portion of the HSD (Huber et al., 2011; Singh et al., 2014). The structure of the SecA-ribosome complex was recently determined at medium resolution (∼11 Å) by cryo-electron microscopy (Singh et al., 2014). However, the molecular mechanism governing the recognition of nascent substrate proteins is not known.

In addition to the catalytic core, most SecA proteins contain a relatively long C-terminal tail (CTT; also known as the C-terminal linker (Hunt et al., 2002)), whose function is not well understood (**figure 1A**). In *E. coli*, the CTT (833-901) contains of a small metal binding domain (MBD; 878-901) and a structurally flexible linker domain (FLD; 833-877). Although it is not required for protein translocation (Fekkes, van der Does, & Driessen, 1997; Or, Boyd, Gon, Beckwith, & Rapoport, 2005; Or, Navon, & Rapoport, 2002), *E. coli* strains producing a C-terminally truncated SecA protein display modest translocation defects (Grabowicz, Yeh, & Silhavy, 2013; Or et al., 2005). The MBD coordinates a Zn^2+^ ion *via* three cysteines and a histidine (or in some species, four cysteines) (Dempsey et al., 2004; Fekkes, de Wit, Boorsma, Friesen, & Driessen, 1999), and in *E. coli* it is required for interaction of SecA with SecB (Breukink et al., 1995; Fekkes et al., 1997; Kimsey, Dagarag, & Kumamoto, 1995). SecB is a molecular chaperone that is required for the secretion of a subset of Sec substrate proteins (Randall & Hardy, 2002). Although not all SecA proteins contain an MBD, the MBD is conserved in many species that do not have a SecB homologue (Dempsey et al., 2004).

**Figure 1.**
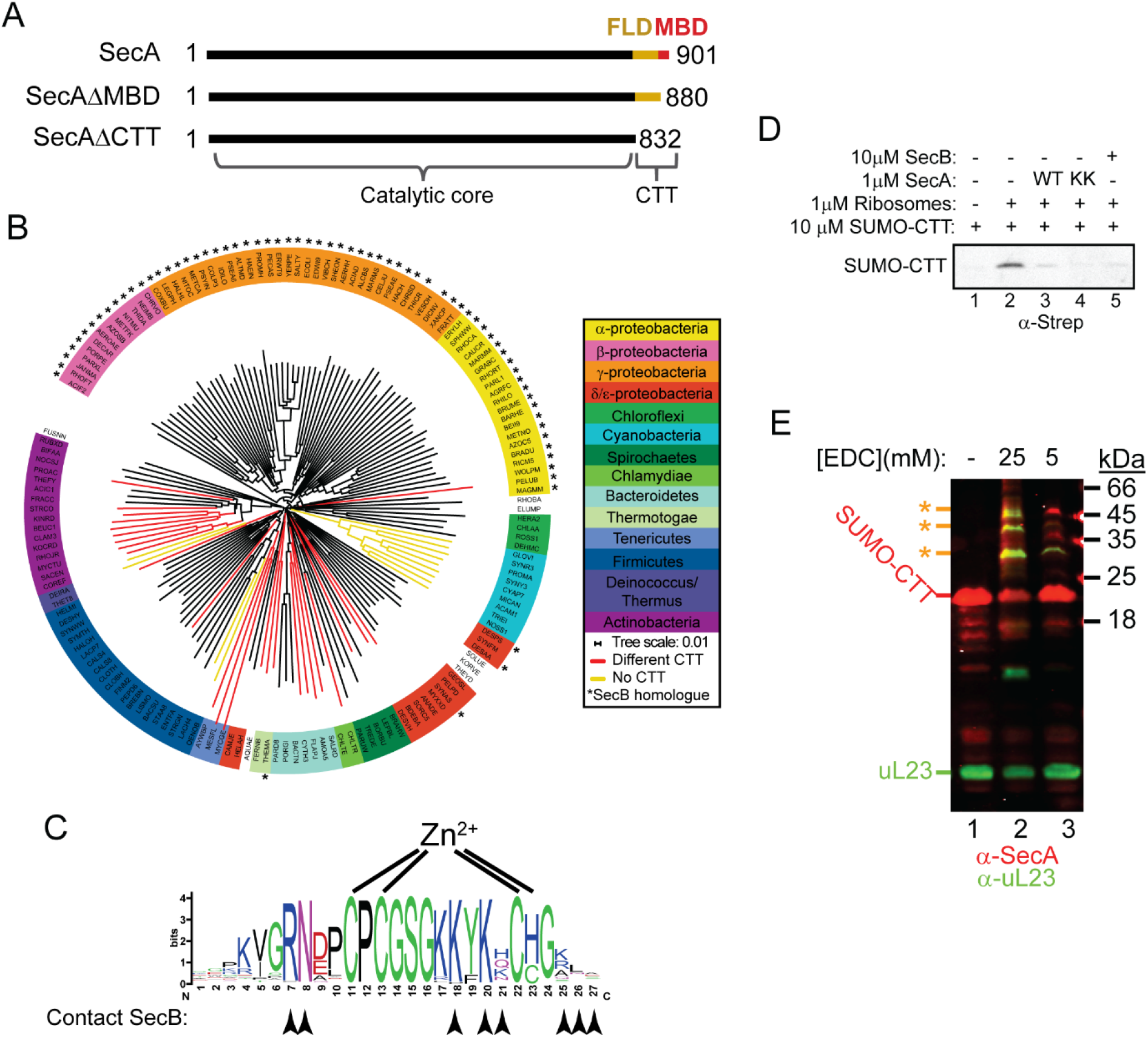
Phylogenetic analysis of the CTT and binding of ***E. coli* CTT to the ribosome.** (A) Schematic diagram of the primary structure of SecA, SecAΔMBD and SecAΔCTT. Structures are oriented with the N-termini to the left, and the amino acid positions of the N-and C-termini are given. Residues of the catalytic core and the CTT are indicated below. Catalytic core, black; FLD, yellow; MBD, red. (B) Phylogenetic tree of the SecA proteins of 156 representative species from 155 different bacterial families. Species names are given as the five-letter organism mnemonic in UniProtKB (The UniProt, 2017). Taxonimical classes are colour-coded according to the legend. Leaves representing SecA proteins with an MBD are coloured black. Those with CTTs lacking a MBD are coloured red, and those that lack a CTT entirely are coloured yellow. Species that also contain a SecB protein are indicated with a star (*). (C) Logo of the consensus sequence of the MBD generated from the 117 species containing the MBD in the phylogenetic analysis. Positions of the metal-coordinating amino acids are indicated above. Amino acids that make contact with SecB in the structure of the MBD-SecB complex (Zhou & Xu, 2003) (1OZB) are indicated by arrowheads below. (D) 10 μM Strep-SUMO-CTT was incubated in the presence or absence of 1 μM ribosomes and layered on a 30% sucrose cushion. Ribosomes were then sedimented through the cushion by ultracentrifugation. Where indicated, 1 μM wild-type SecA, 1 μM containing alanine substitutions at position 625 and 633 (SecA^KK^) or 10 μM tetrameric SecB were included in the binding reaction. Samples were resolved on SDS-PAGE and probed by western blotting using HRP-coupled Streptactin against the Strep tag. (E) 10 μM SUMO-CTT containing an N-terminal Strep(II)-tag was incubated with 1 μM purified ribosomes and treated with 5 mM or 25 mM EDC, as indicated. Samples were resolved by SDS-PAGE and analysed by western blotting by simultaneously probing against SecA (red) and ribosomal protein uL23 (green). Running positions of SUMO-CTT, L23 and crosslinking adducts between them (*) are indicated at left.

In this study, we investigated the role of the CTT in the recognition of nascent substrate proteins by SecA. Phylogenetic analysis of the CTT suggested that it could be involved in binding of SecA to ribosomes, which we confirmed using ribosome cosedimentation and chemical crosslinking approaches. Strikingly, disruption of the MBD alone or the entire CTT had opposing effects on multiple different activities of SecA, suggesting that the CTT affects conformation of the catalytic core. Mass spectrometry, x-ray crystallography, and small-angle x-ray scattering experiments indicated that the FLD is bound in the substrate binding groove and affects the conformation of the PPXD. Finally, site specific chemical crosslinking suggested that binding of the MBD to the ribosome allows full-length SecA to interact with nascent substrate proteins. Taken together, our results provide insight into the molecular mechanism underlying nascent substrate recognition by SecA.

## Results

### Evolutionary distribution of the MBD of SecA

To investigate the evolutionary distribution of the CTT, we analysed the sequences of 156 SecA proteins from bacterial species in 155 phylogenetic families using ClustalOmega (McWilliam et al., 2013). The phylogenetic tree produced by this analysis generally placed SecA proteins from more closely related species (*e.g.* those in the same phylogenetic class) into similar groups (**figure 1B**). The majority of SecA proteins (143) contained a CTT (**figure 1B, red and black branches**). Of these, 117 contained an MBD (**figure 1B, black branches**). A small minority (13) lacked the CTT entirely (**figure 1B, yellow branches**). Of the SecA proteins in the 69 species that contained a SecB homologue (**figure 1B, starred species**), only two lacked an MBD. The strong conservation of the MBD in species that contain SecB suggests that there is strong selective pressure to maintain the MBD in species containing SecB. However, a significant number of species (52) have a SecA protein containing an MBD but lack a SecB homologue. In addition, many of the residues implicated in SecB binding were strongly conserved even in the MBDs of species that lacked SecB (**figure 1C, arrowheads**) (Zhou & Xu, 2003). These results suggested that the MBD has an evolutionarily conserved function in addition to its role in binding to SecB.

### The MBD binds to the ribosome near uL23

Many of the most highly conserved residues in the MBD consensus sequence are positively charged (**figure 1C**), suggesting that the MBD could interact with the negatively charged surface of the ribosome. To determine whether the CTT could bind to ribosomes, we fused the C-terminal 70 amino acids of *E. coli* SecA to the small ubiquitin-like modifier from *Saccharomyces cerevisiae* (SUMO-CTT). Sedimentation of SUMO-CTT through a 30% sucrose cushion during ultracentrifugation was dependent on the presence of ribosomes in the binding reaction, indicating that it could bind to ribosomes (**figure 1D, lanes 1 & 2**). The ribosome binding activity could be attributed to the MBD since a shorter protein fusion containing only the MBD (SUMO-MBD) also cosedimented with ribosomes (**supplemental figure S2**). The presence of purified SecA or SecB in binding reactions inhibited cosedimentation (**figure 1D, lanes 3-5**), indicating that binding to ribosomes and to SecB are mutually exclusive.

We also determined the site of interaction between the CTT and the ribosome using chemical crosslinking. Full-length SecA crosslinks to uL23 in the presence of the non-specific crosslinking agent EDC (Huber et al., 2011). Incubation of SUMO-CTT with ribosomes in the presence of 5 mM and 25 mM EDC resulted in the appearance of several crosslinking products, which crossreacted with antibodies against uL23 and SecA (**figure 1E**). These results suggested that the CTT binds in the vicinity of the opening of the ribosomal exit channel.

### The C-terminal tail affects the affinity of SecA for ribosomes

To determine the contribution of the MBD to binding of SecA to the ribosome, we constructed two C-terminal truncation variants of SecA and measured their affinity for the ribosome using fluorescence anisotropy (Huber et al., 2011) (**figure 1A**). The dissociation constant (K_D_) of the complex between full-length SecA and ribosomes was ∼ 640 nM, similar to previously published figures (Huber et al., 2011). Truncation of the C-terminal 69 amino acids of SecA (SecAΔCTT) caused a modest, but reproducible, increase in the K_D_ of the complex (920 nM) (**figure 2A & table 1**), consistent with the idea that the CTT contributes to ribosome binding. However, truncation of the C-terminal 21 amino acids (SecAΔMBD) significantly increased the affinity of SecA for the ribosome (K_D_ = 160 nM) (**figure 2A & table 1**). These differences in affinity were also evident from the amount of SecA that cosedimented with ribosomes during ultracentrifugation (**figure 2B, lanes 1-6**).

**Figure 2.**
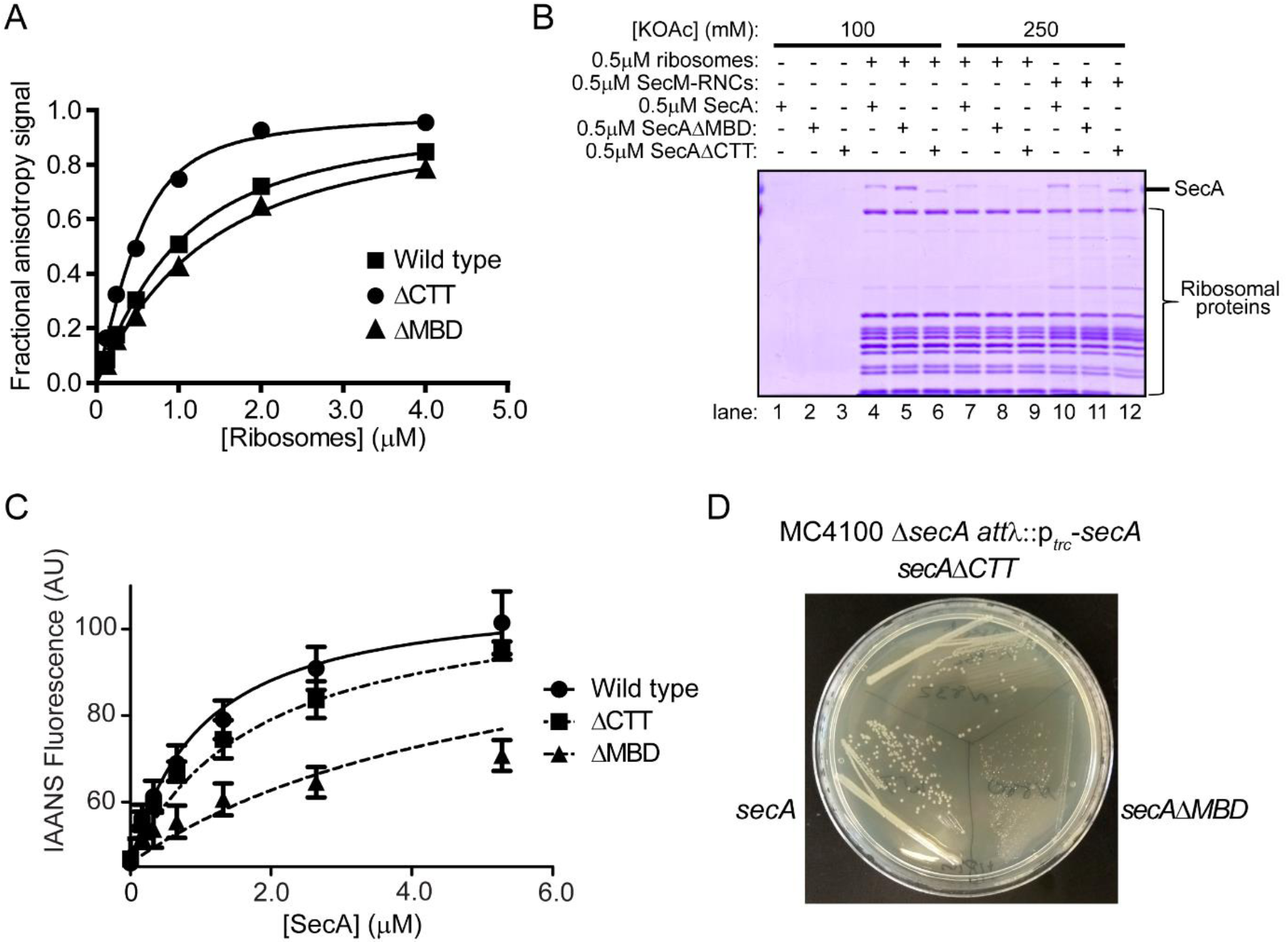
Effect of C-terminal truncations on SecA *in vitro* and *in vivo*. (A) 900 nM Ru(bpy)_2_(dcbpy)-labelled SecA (Wild type; squares), SecAΔMBD (ΔMBD; triangles) or SecAΔCTT (ΔCTT; circles) was incubated in the presence of increasing concentrations of purified 70S ribosomes. The equilibrium dissociation constant (K_D_) of the complex was determined by fitting the increase in fluorescence anisotropy from the Ru(bpy)_2_(dcbpy) (lines; **Table 1**). (B) 0.5 μM SecA, SecAΔMBD or SecAΔCTT was incubated in the absence (lanes 1-3) of ribosomes, in the presence of 0.5 μM vacant 70S ribosomes (lanes 4-9) or in the presence of 0.5 μM RNCs containing nascent SecM peptide (lanes 10-12). Where indicated, binding reactions were incubated in the presence of 100 mM (lanes 1-6) or 250 mM (lanes 7-12) potassium acetate (KOAc). Binding reactions were layered on a 30% sucrose cushion and ribosomes were sedimented through the sucrose cushion by ultracentrifugation. Ribosomal pellets were resolved by SDS-PAGE and stained by Coomassie. (C) 600 nM IAANS-VipB peptide was incubated with increasing concentrations of SecA (Wild type; circles), SecAΔMBD (ΔMBD; triangles) or SecAΔCTT (ΔCTT; squares). The K_D_ for the SecA-peptide complex was determined by fitting the increase in IAANS fluorescence upon binding to SecA (lines; **table 1**). (D) Growth of strains producing SecA (DRH1119; bottom left), SecAΔMBD (DRH1120; bottom right) and SecAΔCTT (DRH1121; top) on LB plates containing 100 μM IPTG.

**Table 1.** Biochemical properties of mutant SecA proteins.

| SecA variant | K<sub>D</sub> Ribosomes<sup>a</sup> | K<sub>D</sub> VipB<sup>b</sup> | Basal ATPase<br>activity<sup>c</sup> | T<sub>M</sub><sup>d</sup> |
| --- | --- | --- | --- | --- |
| Wild type | 640 nM | 0.94 μM | 0.053 ± 0.02 s<sup>-1</sup> | 40.7 ± 0.09 °C |
| SecAΔZnBD | 160 nM | 1.74 μM | < 0.001 s<sup>-1</sup> | 42.0 ± 0.08 °C |
| SecAΔCTT | 920 nM | 5.93 μM | 0.91 ± 0.02 s<sup>-1</sup> | 40.0 ± 0.1 °C |
| SecA<sup>C885A/C887A</sup> | ND<sup>e</sup> | >10 μM | < 0.001 s<sup>-1</sup> | ND<sup>e</sup> |
| SecA<sup>Bpa852</sup> | ND<sup>e</sup> | >10 μM | < 0.001 s<sup>-1</sup> | ND<sup>e</sup> |

<sup>a</sup>Equilibrium dissociation constant of the complex between SecA and non-translating 70S ribosomes as determined by fluorescence anisotropy.
<sup>b</sup>Equilibrium dissociation constant of the complex between SecA and IAANS-labelled VipB peptide as determined by change in fluorescence.
<sup>c</sup>Rate of ATP hydrolysis by SecA in the absence of substrate protein and SecYEG
<sup>d</sup>Denaturation midpoint temperature as determined by the change in circular dichroism at 222 nm.
<sup>e</sup>not determined

### Affinity of truncated SecA proteins for substrate polypeptides

We reasoned that the truncation variants could occupy distinct conformations with differing affinities for the ribosome and that the truncations might therefore affect other activities of SecA. First, we examined binding of SecA to nascent SecM, a SecA substrate protein that has previously been used as a model for binding to nascent chains (Huber et al., 2017; Huber et al., 2011). Binding of SecA to non-translating ribosomes was sensitive to the presence of salt in the binding buffer (**figure 2B, lanes 7-9**). Binding of SecA and SecAΔCTT to ribosome-nascent chain complexes containing nascent SecM (SecM-RNCs) was stable in the presence of 250 mM potassium acetate (**figure 2B, lanes 10 & 12**), indicating that they interacted with nascent SecM. However, nascent SecM did not stabilise binding of SecAΔMBD to RNCs under the same conditions (**figure 2B, lane 11**).

We next examined the effect of the truncations on the affinity of SecA for free substrate protein. To this end, we determined the affinity of SecA for a short peptide (VipB), which was labelled with an environmentally sensitive fluorophore (IAANS-VipB). The amino acid sequence for VipB does not resemble a Sec targeting signal (Pietrosiuk et al., 2011), and binding to VipB was used as a model for binding of SecA to the mature portion of Sec substrate proteins. The affinities of SecA and SecAΔCTT for IAANS-VipB (K_D_ = 0.9 μM and 1.7 μM, respectively) were consistent with the previously reported affinity of SecA for unfolded substrate protein, but the affinity of SecAΔMBD for IAANS-VipB was significantly lower (K_D_ = 5.9 μM) (**figure 2C** & **table 1**) (Gouridis, Karamanou, Gelis, Kalodimos, & Economou, 2009). Furthermore, alanine substitutions in two of the metal-coordinating cysteines (SecA^C885A/C887A^) greatly reduced the affinity of SecA for IAANS-VipB (**table 1**), suggesting that disrupting the structure of the MBD was sufficient to cause this decrease in affinity.

### Effect of alterations to the CTT on the ATPase activity of SecA

To investigate the effect of the truncations on the ATPase activity of SecA, we determined the basal ATPase rates of SecA, SecAΔMBD, SecAΔCTT and SecA^C885A/C887A^. The ATP turnover rate for full-length SecA was 0.05 s^−1^ (**table 1**), consistent with previously reported figures (Huber et al., 2011). Deletion of the entire CTT caused a >10-fold increase in the basal ATPase activity compared to the full-length protein (0.9 s^−1^) (**table 1**), suggesting that the FLD inhibits the ATPase activity of SecA. SecAΔMBD and SecA^C885A/C887A^ did not hydrolyse ATP at a detectable rate, suggesting that the MBD is required to relieve the FLD-mediated autoinhibition.

### C-terminal truncations differentially affect the T_M_ of folding of SecA

To investigate the effect of the C-terminal truncations on the folding of SecA, we examined the secondary structure content and thermal stability of SecA, SecAΔCTT and SecAΔMBD using circular dichroism (CD) spectroscopy. The CD spectra of the three proteins indicated that they were fully folded (**supplemental figure S3A**). However, the denaturation midpoint temperature (T_M_) of SecAΔMBD (42°C) was ∼1.5°C higher relative to that of the full-length protein and ∼2°C higher than that of SecAΔCTT (**table 1, supplemental figure S3B**), indicating that SecAΔMBD was more stably folded than SecA or SecAΔCTT.

### SecA truncation variants have differing abilities to complement the growth defect of a Δ*secA* mutation

To investigate the effect of the C-terminal truncations on the function of SecA *in vivo*, we constructed strains in which the sole copy of the *secA* gene produced a truncated version of the protein in an IPTG-inducible fashion. Because SecA is required for viability, growth of these strains was a reflection of the activity of the SecA variant *in vivo*. Genes producing all three proteins complemented the viability defect of Δ*secA* mutant (**figure 2D**), indicating that they were functional *in vivo*. SecAΔCTT and SecAΔMBD displayed cold-sensitive growth defects compared to the strain expressing wild-type SecA, consistent with a defect in the ability of the proteins to interact with SecB (Shimizu, Nishiyama, & Tokuda, 1997). However, in contrast to cells producing SecAΔCTT, cells producing SecAΔMBD as the sole form of SecA displayed a severe slow growth defect even at the permissive temperature at all IPTG concentrations tested. These results confirmed that the function of SecAΔMBD was also defective *in vivo*.

### The CTT is bound in the substrate binding groove of full-length SecA

In order to affect such a range of activities of SecA, we reasoned that the CTT was likely interacting with the catalytic core of the protein. To investigate this possibility, we incorporated the non-natural amino acid benzophenylalanine (Bpa) into the CTT at positions 852, 893 and 898 using nonsense suppression (Chin, Martin, King, Wang, & Schultz, 2002). Bpa contains a photoactivatable side chain that forms covalent crosslinks to nearby molecules containing C-H bonds. In order to distinguish between early termination products and full-length SecA, we used a variant of SecA containing a biotin-attachment peptide at its C-terminus (SecA-biotin) (Huber et al., 2011; Tagwerker et al., 2006). To aid in purification, we fused SecA to the C-terminus of SUMO, which contained an N-terminal hexahistidine tag. In all cases, the purified protein cross-reacted with streptavidin (**figure 3A**), indicating that Bpa was efficiently incorporated. However, SecA containing Bpa at position 852 (SecA^Bpa852^-biotin) migrated more rapidly than the other proteins in SDS-PAGE (**figure 3A**). Production of this faster migrating species was very efficient: virtually all of the purified protein migrated with a lower apparent molecular weight even before photoactivation (**figure 3B**). In addition, SecA^Bpa852^-biotin exhibited a very low affinity for substrate protein and displayed no detectable ATPase activity (**table 1**). These results suggested that SecA^Bpa852^-biotin contained an internal crosslink, which locked it into a conformation similar to that of SecAΔMBD. To investigate the site of this internal crosslink, we determined the molecular weights of the tryptic peptides of SecA^Bpa852^-biotin using mass spectrometry (MALDI-TOF). As expected, the tryptic peptide containing position 852 (851-877) was absent from the mass spectrum of SecA^Bpa852^-biotin. The only other peptide absent from SecA^Bpa852^-biotin but present in wild-type SecA-biotin was the peptide corresponding to amino acids 361-382 (**figure 3C**), suggesting that the crosslink is to one or more of the residues in this sequence. These amino acids are located in the groove where SecA binds to substrate protein in one of the two strands linking the PPXD to NBD1 (**figure 3D**) (Cranford Smith & Huber, 2018). The site of this crosslink was also consistent with the location of the the extreme N-terminal residues of the FLD in the crystal structure of SecA from *Bacillus subtilis* (1M6N) (Hunt et al., 2002).

**Figure 3.**
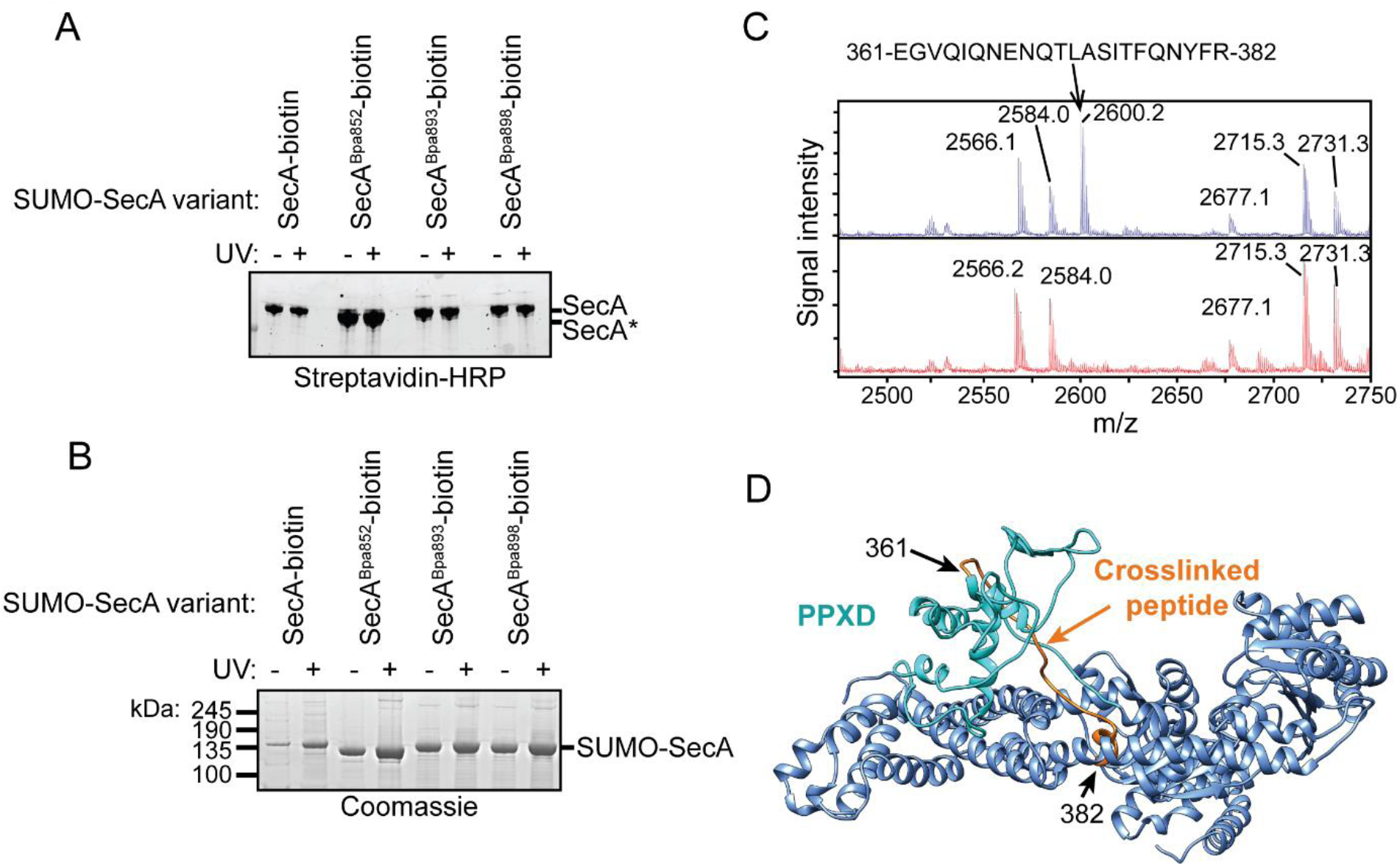
Auto-crosslinking of the CTT to the catalytic core of SecA. (A & B) 1 μM SUMO-tagged SecA-biotin containing Bpa at position 852, 893 or 898 in the CTT was incubated in the absence (-) or presence (+) of light (UV) at 350 nm. The protein samples were resolved using SDS-PAGE and visualised by (A) western blotting against the C-terminal biotin tag or (B) Coomassie staining. The running positions of full-length SUMO-SecA and the internal crosslinking adduct (*) are indicated. (C) Mass spectra of wild-type SecA-biotin (above, blue) and SecA^Bpa852^-biotin (below, red) in the region of 2450-2750 Da region. Wild-type SecA-biotin and SecA^Bpa852^-biotin were exposed to light at 350 nm and subsequently digested with trypsin. The masses of the tryptic fragments were determined using MALDI-TOF. (D) Structure of SecA (2VDA (Gelis et al., 2007)). The main body of the catalytic core is coloured blue, the PPXD is coloured cyan and the tryptic peptide that crosslinks to position 852 (amino acids 361-382) is highlighted in orange. The structural model was rendered using Chimera v. 1.12 (Pettersen et al., 2004).

### Structural analysis of the SecA truncation variants

To investigate the effect of the FLD on the conformation of the SecA catalytic core, we determined the crystal structure of SecAΔMBD at 3.5Å resolution (6GOX; **figure 4A** and **supplemental table S1**). SecAΔMBD crystallised as a symmetric dimer in a head-to-tail configuration (**figure 4A**), similar to that reported for the *E. coli* SecA homodimer in complex with ATP (Papanikolau et al., 2007) (PDB file 2FSG). Consistent with previous studies, the structure of the PPXD was less well defined relative to the other domains of the catalytic core, supporting the idea that the PPXD is structurally mobile (Gold, Whitehouse, Robson, & Collinson, 2013; Zimmer & Rapoport, 2009). However, the FLD was not resolved, and its effect on the structure of the ribosome-binding surface could not be determined.

**Figure 4.**
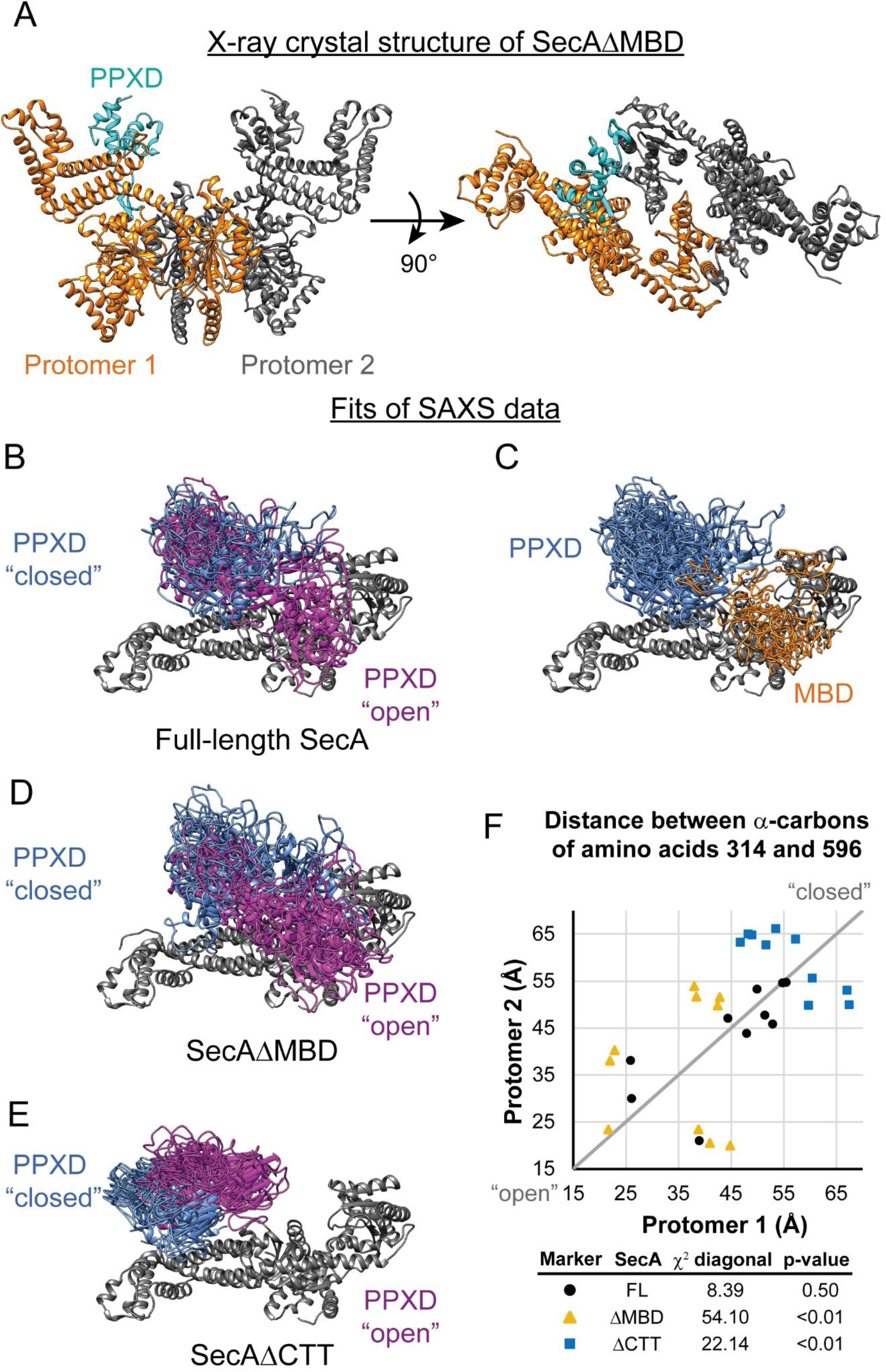
SAXS analysis of SecA truncation variants. (A) X-ray crystal structure of SecAΔMBD at 3.5Å solved by molecular replacement. The asymmetric unit (Protomer 1) is coloured orange with the PPXD highlighted in cyan. The crystallographic mate (Protomer 2) interacts with promoter 1 using an interface similar to that found in 2FSG (Papanikolau et al., 2007). (B-E) Overlay of 10 independent structural models of SecA (B, C), SecAΔMBD (D) and SecAΔCTT (E) generated from fitting to the SAXS data using CORAL. The main body of the catalytic core is coloured grey, and the flexible residues are not displayed. (B, D, E) To facilitate visualization of the asymmetry in the in the dimeric models, both protomeric partners of the dimer were overlaid and the PPXD was coloured (blue/magenta) according to the protomer. The MBD is not displayed in panel B. (C) To facilitate visualization of the position of the MBD in the full-length protein, both protomeric partners of the dimer were overlaid and the MBD of the dimer pair that was located nearest to position 596 of the depicted protomer (orange) was displayed. In panel C, the PPXDs of two protomers, which occupy the same space as the MBDs, are not displayed. (F) Plot of the position of the PPXD in partners of the SecA dimer predicted by structural modelling. The distance between the α-carbon of amino acid 314, which is located near the centroid of the PPXD, and amino acid 596 in NBD2 was determined for each protomer and plotted against the distance in the second protomer. SecA, black circles (FL); SecAΔMBD, orange triangles (ΔMBD); SecAΔCTT, blue squares (ΔCTT). The grey diagonal line indicates the position of the distances if the dimers were symmetric. χ^2^ values to the diagonal were calculated and used to determine p-values to test whether the asymmetry in the dimer was statistically significant.

In order to determine the effect of the truncations on the conformation of the PPXD and the CTT, we investigated the structures of SecA, SecAΔCTT and SecAΔMBD in solution using small-angle x-ray scattering (SAXS) (**supplemental table S2**). The SAXS spectra for all three proteins were similar in the low-*q* region, indicating that the overall shapes of the three proteins were similar, and the radii of gyration suggested that they were dimeric, consistent with previous studies (Woodbury, Hardy, & Randall, 2002) (**supplemental figure S4**). However, the spectra of the three proteins diverged significantly in the mid-*q* region (**supplemental figure S4, arrow**), indicating that there were differences in domain organisation. SecA has been crystallised in several distinct dimer configurations, and the physiological configuration of the dimer and its relevance is an issue of on-going dispute (see discussion in Cranford Smith and Huber (2018)). However, fitting of structural models of the *E. coli* SecA dimer based on PDB files 2FSG (Papanikolau et al., 2007), 2IBM (Zimmer, Li, & Rapoport, 2006), 1M6N (Hunt et al., 2002), 1NL3 (Sharma et al., 2003), 2IPC (Vassylyev et al., 2006) and 6GOX indicated that under the experimental conditions, the conformation of the dimer was similar to that found in 2FSG (χ^2^ = 3.66) and 6GOX (χ^2^ = 5.25) (**supplemental table S3**). To gain insight into the structural differences between the three proteins, we modelled the SAXS data by structural fitting. Initial fitting runs indicated that the CTT was in close proximity to the catalytic core in models of full-length SecA and SecAΔMBD, and further fitting runs were carried out by fixing the FLD to a position consistent with the Bpa crosslinking results. These models suggested that the PPXD was positioned much closer to NBD2 (i.e. more “open”) in SecA and SecAΔMBD than in SecAΔCTT (p = 2.0 × 10^−5^ and 1.1 × 10^−7^, respectively) (**figure 4F**). In models of SecAΔMBD and SecAΔCTT, the PPXDs in the two protomers of the dimers were positioned asymmetrically (p = 1.8 × 10^−8^ and 0.0085, respectively) (**figure 4B-D, F**). Finally, in models of full-length SecA, the MBD was positioned between NBD2 and the C-terminal portion of the HSD (amino acids 756-832) in both protomers of the dimer (**figure 4E**). Localisation of the MBD to this region would position it directly adjacent to the ribosome-binding surface on the catalytic core (Huber et al., 2011; Singh et al., 2014).

### Site specific crosslinking of SecA to ribosomes

To investigate the effect of the CTT on binding of SecA to the ribosome, we incorporated Bpa at 12 positions located on the face of SecA that binds to the ribosome, including positions 56, 260, 299, 399, 406, 625, 647, 665, 685, 695, 748 and 796 (**figure 5A & B**) (Huber et al., 2011; Singh et al., 2014). In the presence of purified 70S ribosomes, SecA containing Bpa at positions 399 (SecA^Bpa399^) and 406 (SecA^Bpa406^) produced an additional high molecular weight band in SDS-PAGE, which was recognised by α-SecA antiserum (**figure 5C**). Analysis of the high-molecular weight band produced by SecA^Bpa399^ by mass-spectrometry (LC-MS/MS) indicated that it was an adduct between SecA and ribosomal protein uL29. uL29 is located adjacent to uL23 on the ribosomal surface, and both F399 and K406 appear to contact uL29 in the structure of the SecA-ribosome complex determined by cryo-electron microscopy (**figure 5A**) (Singh et al., 2014). To investigate the effect of a nascent chain on crosslinking between SecA^Bpa399^ and uL29, we incubated SecA^Bpa399^ with RNCs containing nascent SecM or maltose binding protein (MBP) (**figure 5D** and **supplemental figure S5**). The presence of a nascent polypeptide completely inhibited crosslinking of SecA^Bpa399^ to uL29, suggesting that binding to nascent polypeptide causes a conformational change in SecA that affects its interaction with the ribosomal surface. However, SecAΔMBD^Bpa399^ produced crosslinks to uL29 in both the presence and the absence of nascent polypeptide (**figure 5D**), suggesting that displacement of the FLD is required for interaction with nascent substrate proteins.

**Figure 5.**
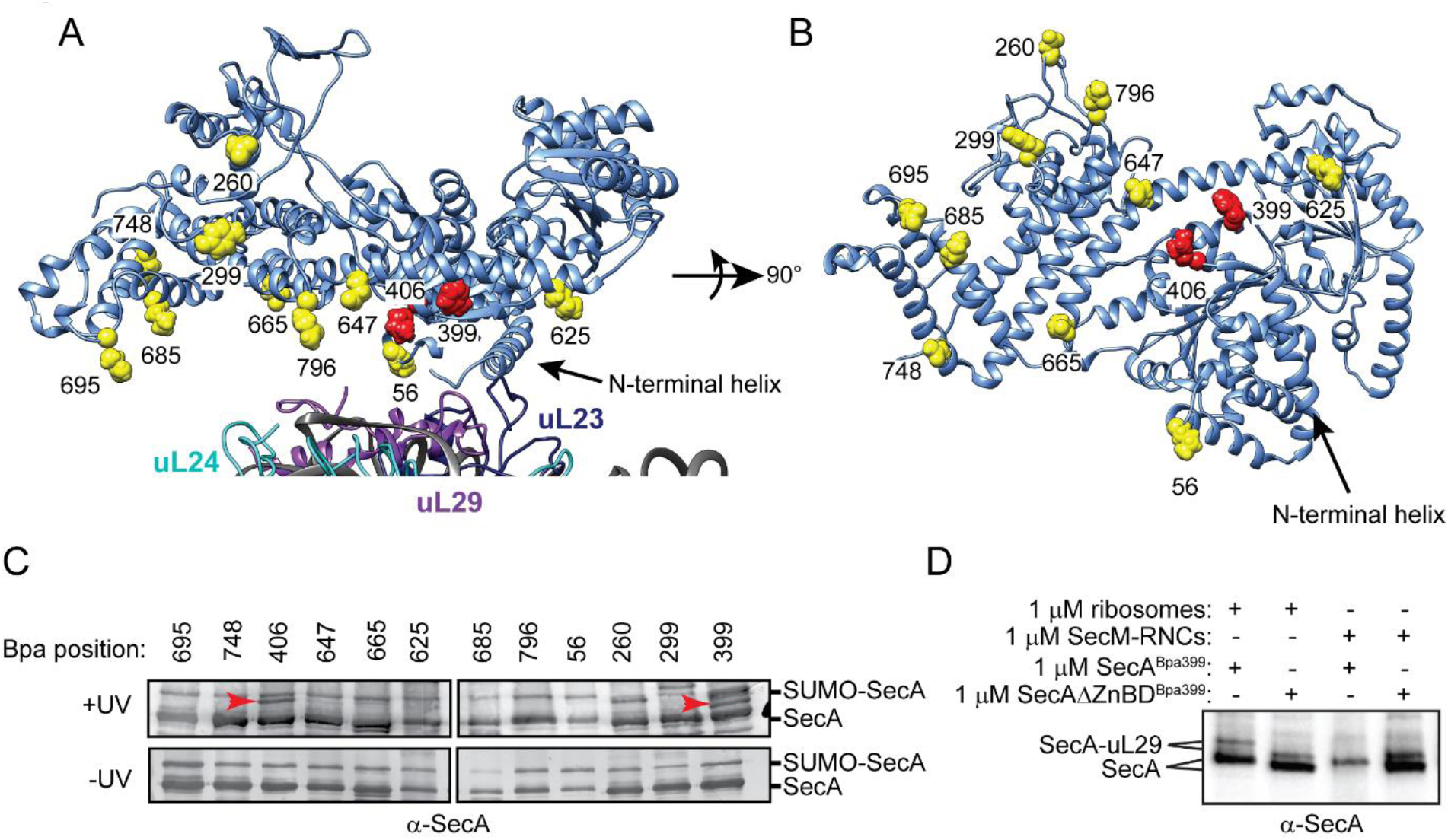
Site specific crosslinking of SecA to purified ribosomes and ribosome-nascent chain complexes. (A & B) Sites of incorporation of Bpa in the structure of *E. coli* SecA. (A) Fit of the high resolution structure of SecA (PDB code 2VDA (Gelis et al., 2007)) and the 70S ribosome (PDB code 4V4Q (Schuwirth et al., 2005)) to the cryoEM structure of the SecA ribosome complex (EMD-2565 (Singh et al., 2014)). (B) View of SecA from the ribosome-interaction surface. Amino acid positions where Bpa was incorporated are represented in space fill (yellow). Positions that crosslink to ribosomal proteins are coloured red. The locations of the N-terminal α-helix of SecA and of ribosomal proteins uL23 (dark blue), uL29 (purple) and uL24 (cyan) are indicated. Structural models were rendered using Chimera v. 1.12 (Pettersen et al., 2004). (C) Bpa-mediated photocrosslinking of SecA variants to vacant 70S ribosomes. 1 μM purified ribosomes were incubated with 1 μM SecA containing BpA at the indicated position and exposed to light at 350 nm (above) or incubated in the dark. Crosslinking adducts consistent with the molecular weight of a covalent crosslink to ribosomal proteins are indicated with red arrowheads. The positions of full-length SecA and uncleaved SUMO-SecA protein are indicated to the right. (D) 1 μM SecA^Bpa399^ or SecAΔMBD^Bpa399^ was incubated with 1 μM non-translating 70S ribosomes or 1 μM arrested RNCs containing nascent SecM (SecM-RNCs) and exposed to light at 350 nm. The positions of full-length SecA and the SecA-uL29 crosslinking adduct are indicated. In C & D, samples were resolved using SDS-PAGE and probed by western blotting using anti-SecA antiserum.

## Discussion

Our results suggest that the CTT controls the conformation of SecA and regulates its activity. Disruption of the MBD (i) increases the affinity of SecA for the ribosome, (ii) decreases the affinity of SecA for substrate protein, (iii) inhibits the ATPase activity of SecA, (iv) increases the thermal stability of SecA, (v) prevents SecA from undergoing a conformational change upon binding to nascent substrate protein and (vi) causes a defect in SecA function *in vivo*. However, SecAΔCTT behaves very similarly to full-length SecA, indicating that the FLD mediates these effects. Chemical crosslinking and structural modelling of the SAXS data for wild-type SecA suggest that the FLD causes these effects by binding in the substrate protein binding groove and causing a conformational change in the PPXD. Gold et al. (2013) have suggested that opening of the PPXD when SecA is bound to substrate protein (*i.e.* enclosing the substrate protein in the binding groove by the PPXD clamp) activates the ATPase activity of SecA. Our work suggests that enclosure of the FLD by the PPXD has the opposite effect—that is, autoinhibiting SecA. We propose that the MBD is the key for unlocking this autoinhibited conformation in the full-length protein. Previous work suggests that interaction of the MBD with SecB increases the affinity of SecA for polypeptides (Gelis et al., 2007). Our results indicate that the MBD also binds to the ribosome, and modelling of the SAXS data reveal that the MBD is ideally positioned to interact with the ribosomal surface (Singh et al., 2014).

Taken together, our results allow us to propose a putative mechanism for the recognition of nascent substrate proteins by SecA (**figure 6**): (i) interaction of the MBD with the ribosomal surface upon binding of SecA to the ribosome destabilises binding of FLD in the substrate protein binding groove; (ii) destabilisation of the FLD allows SecA to sample nascent polypeptides; (iii) the stable interaction of SecA with nascent substrate protein displaces the FLD from the substrate binding groove; (iv) binding of SecA to nascent substrate protein causes a conformational change in SecA that leads to release from the ribosome.

**Figure 6.**
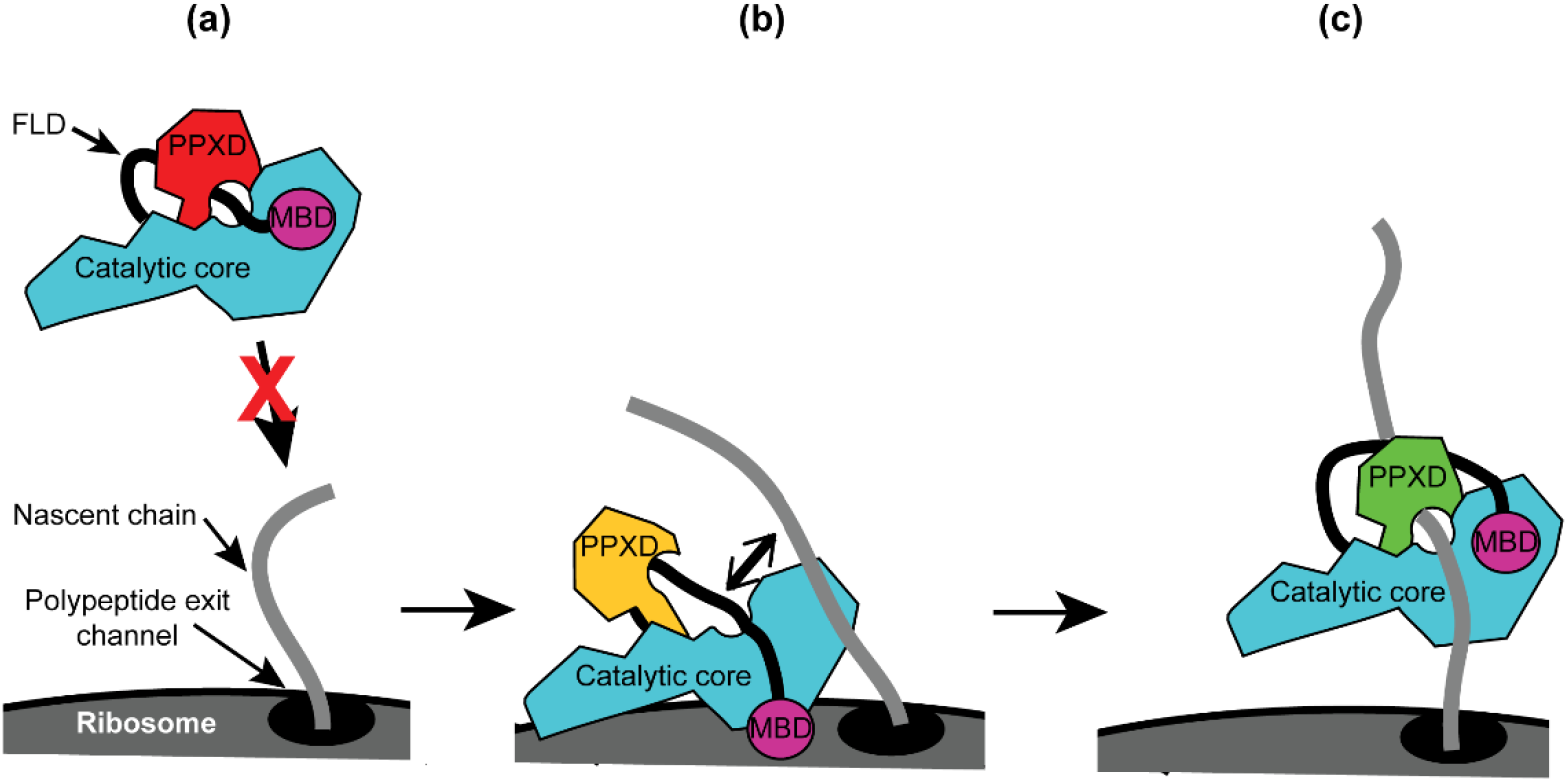
Diagram of the proposed mechanism for recognition of nascent substrate proteins by SecA. (a) In solution, SecA occupies an autoinhibited conformation with the FLD bound stably in the substrate protein binding site and the PPXD in the open conformation. (b) Binding of both the catalytic core and the MBD to the ribosomal surface causes the PPXD to shift to the open conformation, which destabilises binding of the FLD and allows SecA to sample nascent polypeptides. (c) Binding to the nascent substrate protein displaces the FLD from the substrate protein binding site and the PPXD returns to the open conformation, stabilising this interaction. Binding to nascent substrate releases SecA from the ribosomal surface.

While our results clearly indicate that the CTT is responsible for autoinhibition of SecA, the physiological role of this autoinhibition is not clear. One possibility is that it prevents the spurious interaction of SecA with non-substrate proteins in the cytoplasm. Restricting the ability of SecA to interact with unfolded polypeptides to certain conditions (*e.g.* in the presence of the ribosome or SecB) would effectively reduce the pool of potential substrate proteins that SecA needs to screen. Indeed, artificially increasing the pool of SecA-substrate proteins causes translocation defects *in vivo* (Muller, Reinert, & Malke, 1989; Oliver & Beckwith, 1982; Wagner et al., 2007). In addition, the spurious translocation of certain proteins (*e.g.* β-galactosidase) could be toxic (Emr, Schwartz, & Silhavy, 1978; van Stelten, Silva, Belin, & Silhavy, 2009).

While the basic features of the catalytic core of SecA are highly conserved amongst bacteria, the SecA proteins of different phylogenetic groups contain a diverse array of loops and extensions. For example, our phylogenetic analysis indicated that a large number of bacterial species contain alternative CTTs with structures that are significantly different from that of *E. coli*. These differences could allow the activity of SecA to be regulated in response to interaction with a different subset of interaction partners. Nonetheless, many of these alternative CTTs are highly positively charged (*e.g.* those of many Actinobacteria), suggesting that they may retain the interaction with the ribosome. Some phylogenetic groups (*e.g.* the Cyanobacteria) lack a CTT entirely. However, most of these species contain large loops in between the conserved elements of the catalytic core of the protein. Indeed, *E. coli* SecA also contains a “variable” subdomain in NBD2 (amino acids ∼519-547), which has been proposed to regulate its activity (Das et al., 2012). It is therefore reasonable to speculate that large loops in between the conserved features of the catalytic core might carry out functions analogous to the CTT in *E. coli*.

## Methods

### Chemicals and media

All chemicals were purchased from Sigma-Aldrich unless otherwise indicated. Rabbit anti-SecA antiserum was a laboratory stock. Sheep anti-uL23 antiserum was a kind gift from R. Brimacombe. IR700-labelled anti-rabbit and IR800-labelled anti-sheep antibodies were purchased from Rockland (Philadelphia, PA). Horseradish peroxidase (HRP)-labelled anti-rabbit antibody was purchased from GE Healthcare. HRP-coupled streptactin was purchased from IBA Lifesciences (Goettingen, Germany). Bpa was purchased from Bachem (Santa Cruz, CA). Strains were grown in lysogeny broth (LB) containing kanamycin (30 μg/ml) or ampicillin (100 μg/ml) as required.

### Strains and plasmids

Strains and plasmids were constructed using standard methods (Miller, 1992; Sambrook & Russell, 2001). For protein-expression plasmids, the DNA encoding full-length SecA or fragments of SecA were amplified by PCR and ligated into plasmid pCA528 (His_6_-SUMO) or pCA597 (Strep_3_-SUMO) using the *BsaI* and *BamHI* restriction sites (Andreasson, Fiaux, Rampelt, Mayer, & Bukau, 2008). UAG stop codons were introduced into plasmids expressing His-SUMO-SecA or His-SUMO-SecA-biotin using QuikChange (Agilent). Strains DRH1119, DRH1120 and DRH1121, which produce were constructed as follows: *secA* genes producing full-length SecA, SecAΔMBD and SecAΔCTT were cloned into pDSW204 (Weiss, Chen, Ghigo, Boyd, & Beckwith, 1999) and introduced onto the chromosome of strain DRH663 (MC4100 Δ*secA*::Kan^R^ + pTrcSpc-*secA*)(Huber et al., 2011) using lambda InCh (Boyd, Weiss, Chen, & Beckwith, 2000). The SecA-producing plasmid was then cured from these strains by plating on LB containing 1 mM isopropyl-thiogalactoside (IPTG). All three strains required > 10μM IPTG for growth on LB.

### Phylogenetic analysis

The sequences of SecA for the given UniProtKB entry names (The UniProt, 2017) were analysed using ClustalOmega (McWilliam et al., 2013). The unrooted phylogenetic tree was rendered using iTOL (Ciccarelli et al., 2006). The logo of the consensus MBD sequence was generated using WebLogo (https://weblogo.berkeley.edu/logo.cgi).

### Ribosome and protein purification

Ribosomes and arrested RNCs were purified as previously described (Huber et al., 2011; Rutkowska et al., 2008). For all SecA proteins, BL21(DE3) (laboratory stock) was transformed with the appropriate plasmid and grown in LB in the presence of kanamycin at 37°C to OD_600_ ∼1, induced using 1 mM IPTG and shifted to 18°C overnight. Cells were then harvested by centrifugation and lysed by cell disruption in buffer (50 mM K·HEPES, pH 7.5, 500 mM NaCl and 0.5 mM TCEP [tris(2-carboxyethyl)phosphine]) containing cOmplete EDTA-free protease inhibitor cocktail (Roche). Unlabelled His-tagged proteins were affinity purified by passing over a 5 ml Ni-NTA HiTrap column (GE Healthcare), washed with buffer containing 50 mM imidazole and eluted from the column in buffer containing 250 mM imidazole. The eluted protein was cut with the SUMO-protease Ulp1 and the cut SUMO was removed by passing over a 5 ml Ni-NTA HiTrap column. The partially purified protein was then concentrated (Centricon) and purified by size exclusion chromatography using a sepharose S-200 column (GE Healthcare). Bpa-labelled proteins were purified as described by Huber et al. (2017). For Strep-tagged proteins, lysates from cells producing SUMO-CTT and SUMO-MBD were passed over streptactin-coupled sepharose beads (IBA Lifesciences), washed extensively with buffer and eluted using buffer containing 10 mM desthiobiotin.

### Western blotting

Western blots were carried out as previously described (Sambrook & Russell, 2001). Protein samples were resolved using “Any kD” SDS-PAGE gels (BioRad) and transferred to nitrocellulose membranes. Membranes were probed using the indicated primary and secondary antisera or with HRP-streptactin. For HRP-based detection, membranes were developed using ECL (GE Healthcare) and visualised using a BioRad Gel-Doc. For IR700-and IR800-based detection, membranes were visualised using a LI-COR Odyssey scanner.

### Ribosome cosedimentation

Ribosome cosedimentation experiments were carried out as previously described (Huber et al., 2017). Binding reactions were incubated in buffer containing 10 mM HEPES potassium salt, pH 7.5, 100 mM potassium acetate, 10 mM magnesium acetate, 1 mM β-mercaptoethanol for >10 minutes. The reaction mixture was then layered on top of a 30% sucrose cushion made with the same buffer and centrifuged at >200,000 x *g* for 90 minutes. The supernatant was discarded. The concentration of ribosomes in the pellet fractions were normalised using the absorbance at 260 nm.

### Fluorescence anisotropy

Determination of the K_D_ of the SecA-ribosome complex by fluorescence anisotropy was determined as previously described (Huber et al., 2011). SecA, SecAΔMBD and SecAΔMBD were labelled with Ru(bpy)_2_(dcbpy) and the fluorescence anisotropy was determined on a Jasco FP-6500 fluorometer containing an ADP303 attachment using an excitation wavelength of 426 nm (slit width 5 nm) and an emission wavelength of 640 nm (slit width 10 nm).

### CD spectroscopy

The CD spectra of 2µM solutions of full-length SecA, SecAΔMBD, or SecAΔCTT in 10 mM potassium phosphate buffer (pH 7.5) were measured at temperatures that promote folding (10°C) and denaturation (85°C) in a 0.1cm cuvette using a Jasco J750 CD spectrometer. For thermal titrations, the temperature was raised 0.5 K/min from 20 to 60°C and circular dichroism was measured at 222 nm.

### Peptide binding

600 nM VipB peptide labelled with IAANS (Pietrosiuk et al., 2011) was incubated with increasing concentrations of SecA or the indicated SecA variant. The increase in IAANS fluorescence upon binding of SecA was measured using a Jasco FP-6500 fluorometer or a BMG Labtech CLARIOStar.

### ATPase assays

ATPase activities were determined by measuring the rate of NADH oxidation in a coupled reaction (Kiianitsa, Solinger, & Heyer, 2003). 1μM SecA, or the respective SecA variant, was added to a solution containing 250 mM NADH, 0.5 mM phosphoenolpyruvate, 2 mM ATP, 20/ml lactate dehydrogenase, 100 U/ml pyruvate kinase and incubated at 25°C, 50 mM K·HEPES, pH 7.5 and 500 mM NaCl. The decrease in absorbance at 340 nm from the oxidation of NADH to NAD^+^ was measured using an Anthos Zenyth 340rt (Biochrom) absorbance photometer equipped with ADAP software. The rate of ATP hydrolysis was determined from the rate of NADH oxidation by dividing the rate of decrease in the absorbance by the extinction coefficient for NADH (6220 M^−1^ cm^−1^).

### Chemical crosslinking

Photo-crosslinking of Bpa SecA and non-specific crosslinking of SUMO-CTT to the ribosome using 1-ethyl-3-(3-dimethylaminopropyl) carbodiimide (EDC) were carried out as described previously (Huber et al., 2017; Huber et al., 2011).

### Mass spectrometry

Auto-crosslinked samples were digested with sequencing grade trypsin and the masses of the tryptic peptides were determined using MALDI-TOF mass spectrometry. The identity of ribosomal crosslinking adducts was determined by excising the protein band from a Coomassie-stained gel and analysing the tryptic peptide fragments using liquid chromatography-tandem mass spectrometry (LC-MS/MS) identification (The Advanced Mass Spectrometry Facility, School of Biosciences, University of Birmingham).

### X-ray crystallography

SecAΔMBD was crystallised by mixing 2 μl of purified protein (100 μM) with 2 μl of a 9:1 mixture of Peg Ion 2 # 39 buffer (0.04M Citric acid, 0.06M BIS-TRIS propane, pH6.4, 20% Polyethylene glycol 3,350) and Morpheus condition 47, box 2 (0.1M Amino acids, 0.1M Buffer system 3, pH 8.5, 50% v/v Precipitant Mix 3) in a 48-well MRC MAXI plate (Molecular Dimensions, Newmarket, UK). Crystals appeared within six days and were fully matured by two weeks. Crystals were analysed at Diamond light source, and the structure was solved by molecular replacement using PDB file 2FSG at 3.5 Å resolution (**Table S1**). The structure was deposited at RCSB under PDB file 6GOX.

### SAXS measurements

Synchrotron radiation X-ray scattering data were collected on the ESRF BM29 BioSAXS beamline (Grenoble) (**Table S2**). An in-line Superose 6 column was used to ensure that the protein was free from aggregates and that it occupied a single oligomeric state during data collection. The sample-to-detector distance was 3 m, covering a range of momentum transfer s = 0.03-0.494 Å^−1^ (s = (4π·sin θ)/ λ, where 2θ is the scattering angle, and λ = 0.992 Å is the X-ray wavelength). Data from the detector were normalized to the transmitted beam intensity, averaged, placed on absolute scale relative to water and the scattering of buffer solutions subtracted. All data manipulations were performed using PRIMUS*qt* and ATSAS (Petoukhov et al., 2012). The forward scattering *I*(*0*) and radius of gyration, *R*_*g*_ were determined by Guinier analysis. These parameters were also estimated from the full scattering curves using the indirect Fourier transform method implemented in the program GNOM, along with the distance distribution function *p*(*r*) and the maximum particle dimensions *D*_*max*_. Molecular masses of solutes were estimated from SAXS data by comparing the extrapolated forward scattering with that of a reference solution of bovine serum albumin. Computation of theoretical scattering intensities was performed using the program CRYSOL.

### Molecular modelling of SAXS data

SAXS data has been deposited at the SASBDB (www.sasbdb.org) with accession codes: SASDDY9 (full-length SecA), SASDDZ9 (SecAΔMBD) and SASDE22 (SecAΔCTT). For modelling based on SAXS data, multiple runs were performed to verify the stability of the solution, and to establish the most typical 3D reconstructions using DAMAVER. Guinier analysis of the SAXS data indicated that the protein was dimeric under the conditions used for SAXS. Structural models of the *E. coli* SecA dimer were generated by aligning the structure of SecA in PDB file 2VDA to PDB files 2FSG, 2IBM, 2IPC, 1M6N, 1NL3 and 6GOX using PyMol v. 1.8.0.5 and refining using GROMACS (Pronk et al., 2013). Because the CTT is not resolved in the structures used for modelling, these models were fit to the SAXS spectrum of SecAΔCTT using FoXS (Schneidman-Duhovny, Hammel, Tainer, & Sali, 2016) (**supplemental table S3**). The structures of full-length SecA, SecAΔMBD and SecAΔCTT were modelled by fitting the 2FSG dimer to the respective SAXS data by multi-step rigid body refinement using CORAL (Petoukhov et al., 2012). The positions of NBD1, NBD2, HSD and HWD were fixed in all models. The regions corresponding to residues 1-8, 220-231, 367-375 were defined as linkers and modelled as flexible. The PPXD was allowed rigid body movement in all three models. The FLD (residues 829-832 in SecAΔCTT and 829-880 in SecAΔMBD and full-length SecA) were modelled as flexible. For full length SecA, the MBD was modelled using PDB file 1SX0 and allowed rigid-body movement. Because initial modelling indicated that the FLD was in close contact with the catalytic core, and because photocrosslinking indicated the FLD was bound in the substrate binding groove, residues 851-854 were modelled to form a small β-sheet with residues 222-225 and 373-375 and allowed rigid body movement. All ten independently generated fits for SecA and SecAΔMBD produced plausible structural models (χ^2^ = 0.97 ± 0.02 and 1.87 ± 0.09, respectively). Ten of 15 of the fits of SecAΔCTT produced plausible structural models (χ^2^ = 1.82 ± 0.18). In the remaining five fits, the position of the PPXD in one of the two protomers was inconsistent with previously published structures of SecA and occupied a non-realisitic conformation, suggesting increased mobility of the PPXD in SecAΔCTT.

## Acknowledgements

We thank B. Zachmann-Brand, Jingli Yu and the functional genomic facility (School of Biosciences, University of Birmingham) for technical assistance. We thank A. McNally and members of the Bukau, Mayer, Henderson, Lund, Grainger, Cole and Rossiter groups for helpful advice and discussions.

---

**Supplemental figure S1.**
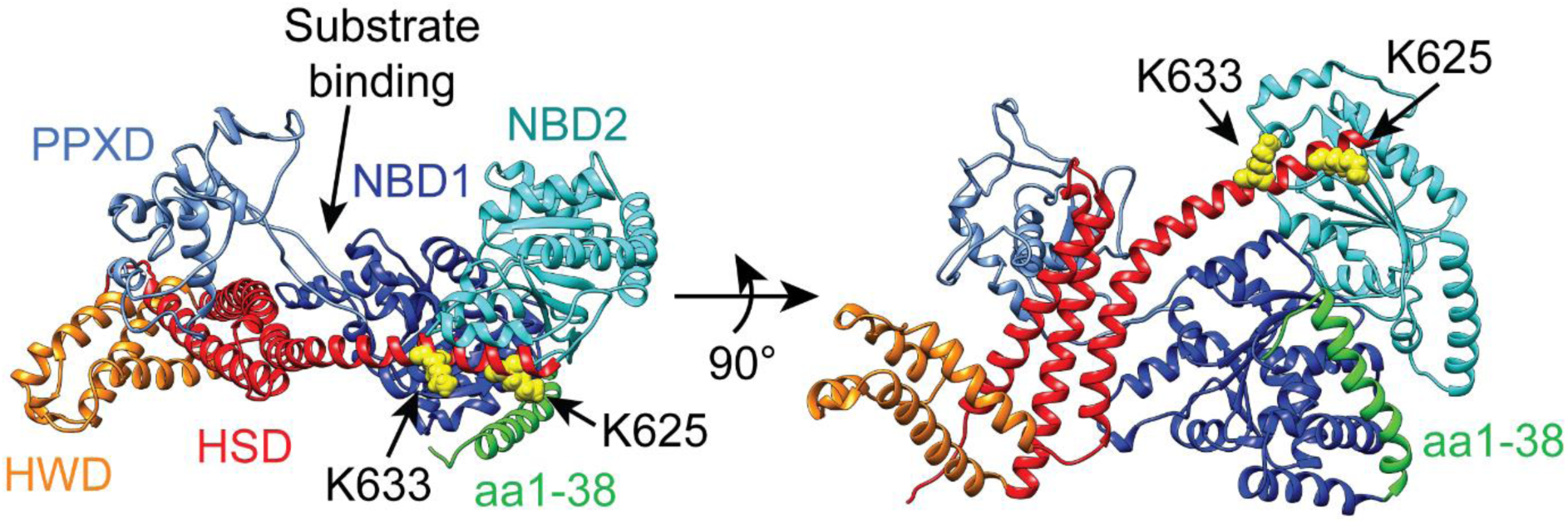
Structural model of the catalytic core of SecA in the “closed” conformation. Structural model of *E. coli* SecA from PDB file 2VDA (Gelis et al., 2007) in ribbon diagram. The model is coloured according to domains described in the main text. NBD1, dark blue; NBD2 cyan; PPXD, light blue; HSD, red; HWD, orange. The side chains of lysines 625 (K625) and 633 (K633), which were identified by Huber et al. (2011) to be involved in ribosome binding, are depicted in space-fill. The N-terminal α-helix (aa1-38), which was identified by Singh et al. (2014) to be involved in ribosome binding is coloured green. The CTT is not resolvable in high-resolution structures of SecA and is therefore not depicted.

**Supplemental figure S2 (associated with figure 1).**
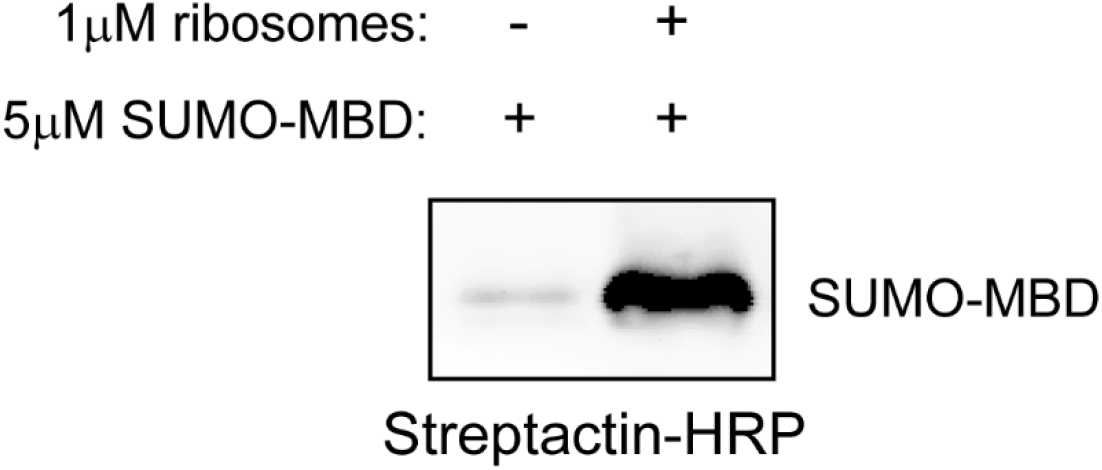
SUMO-MBD cosediments with ribosomes. The C-terminal 27 amino acids of SecA were fused to the C-terminus of Strep-tagged SUMO (SUMO-MBD). 5 μM SUMO-MBD was incubated in the presence or absence of 1 μM 70S ribosomes. Binding reactions were layered on a 30% sucrose cushion and subjected to ultracentrifugation to sediment ribosomes. The pellet fractions were then resolved by SDS-PAGE and analysed by western blotting against the Strep tag.

**Supplemental figure S3 (associated with table 1).**
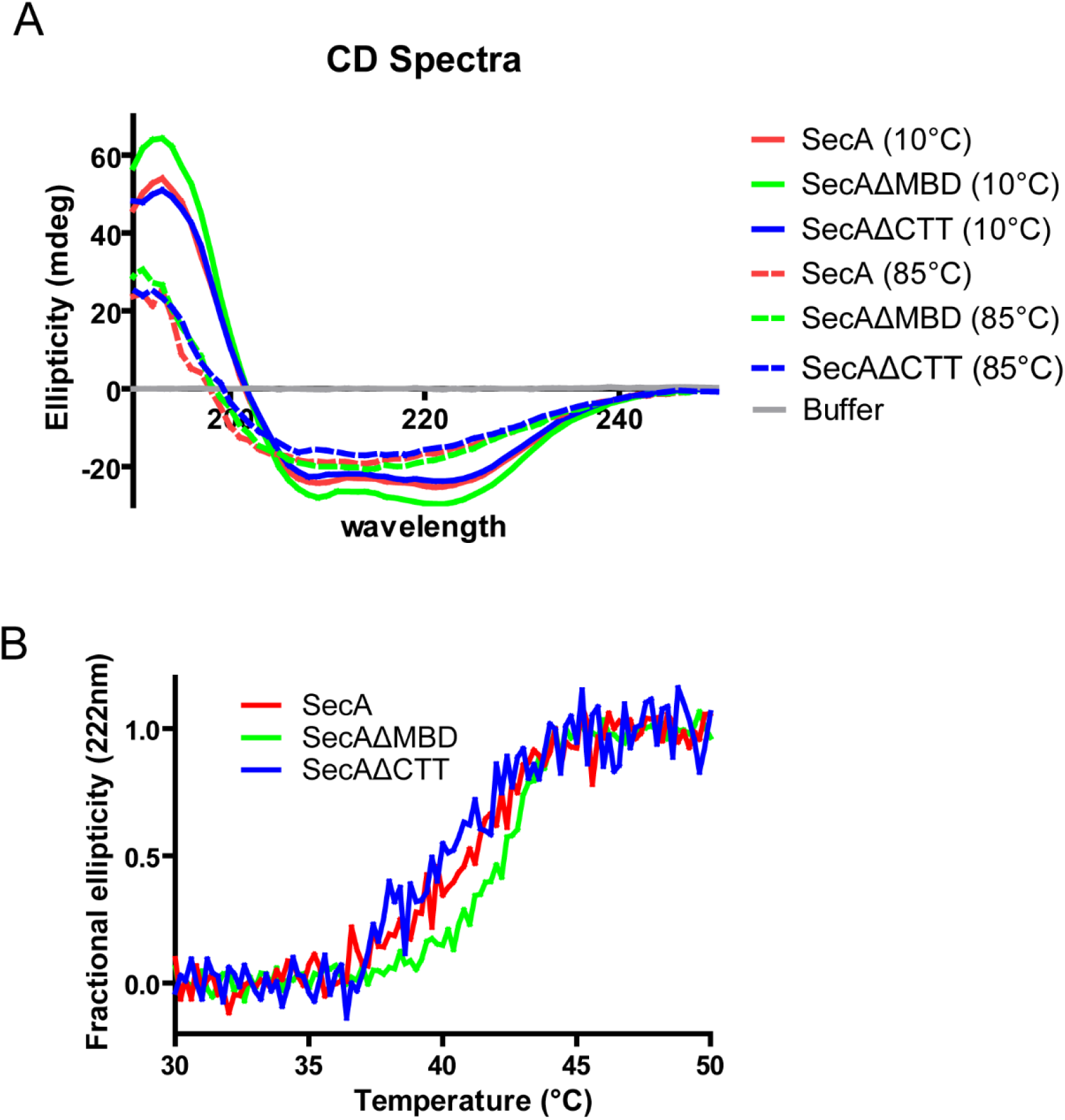
CD spectra and thermal denaturation plots of SecA, SecAΔMBD and SecAΔCTT. (A) Far-UV circular dichroism (CD) spectra of 2 µM solutions of SecA, SecAΔMBD, and SecAΔCTT in 10mM potassium phosphate (pH 7.5). (B) Representative plot of the thermal denaturation of SecA as determined by CD spectroscopy. The α-helical content of 2 µM solutions of SecA, SecAΔMBD, and SecAΔCTT in 10mM potassium phosphate (pH 7.5) was determined by measuring molar ellipticity at 222 nm while the temperature of the solution was raised from 30 to 50°C. The T_M_s given in **Table 1** were determined by van’t Hoff analysis.

**Supplemental table S1.** Data collection and refinement statistics for the crystal structure of SecAΔMBD.

|  | **SecAΔMBD** |
| --- | --- |
| **Wavelength** | 0.9795 |
| **Resolution range** | 47.5 - 3.5 (3.625 - 3.5)* |
| **Space group** | C 1 2 1 |
| **Unit cell** | 163.1 106.8 76.0 90 102.8 90 |
| **Total reflections** | 32115 (3197)* |
| **Unique reflections** | 16106 (1609)* |
| **Multiplicity** | 2.0 (2.0)* |
| **Completeness (%)** | 99.66 (99.81)* |
| **Mean I/sigma(I)** | 13.51 (1.51)* |
| **Wilson B-factor** | 132.20 |
| **R-merge** | 0.03651 (0.5532)* |
| **R-meas** | 0.05163 (0.7823)* |
| **R-pim** | 0.03651 (0.5532)* |
| **CC1/2** | 1 (0.672)* |
| **CC\*** | 1 (0.897)* |
| **Reflections used in refinement** | 16106 (1609)* |
| **Reflections used for R-free** | 785 (89)* |
| **R-work** | 0.2316 (0.3407)* |
| **R-free** | 0.2927 (0.3891)* |
| **CC(work)** | 0.900 (0.645)* |
| **CC(free)** | 0.869 (0.405)* |
| **No. non-hydrogen atoms** | 6182 |
| **Protein residues** | 777 |
| **RMS(bonds/angles)** | 0.011/1.47 |
| **Ramachandran (%)** | 91.42/5.07/3.51 |
| favoured/allowed/outliers |  |
| **Rotamer outliers (%)** | 3.64 |
| **Clashscore** | 14.32 |
| **Average B-factor** | 158.55 |
| **PDB** | 6GOX |

\*Statistics for the highest-resolution shell are shown in parentheses.

**Supplemental figure S4 (associated with figure 4).**
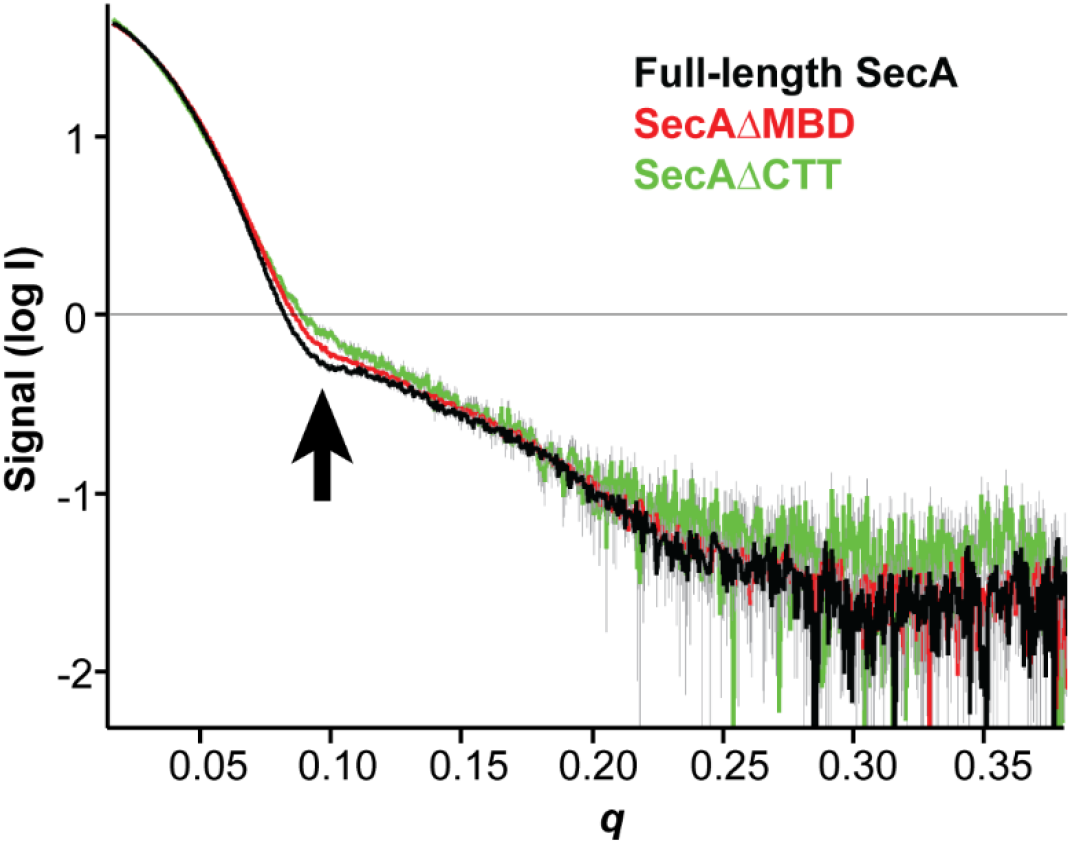
SAXS analysis of the solution structure of SecA, SecAΔMBD and SecAΔCTT. X-ray scattering plots for SecA (black), SecAΔMBD (red) and SecAΔCTT (green). The region of divergence between the three SAXS traces in the mid-*q* region is indicated (black arrow).

**Supplemental table S2.**
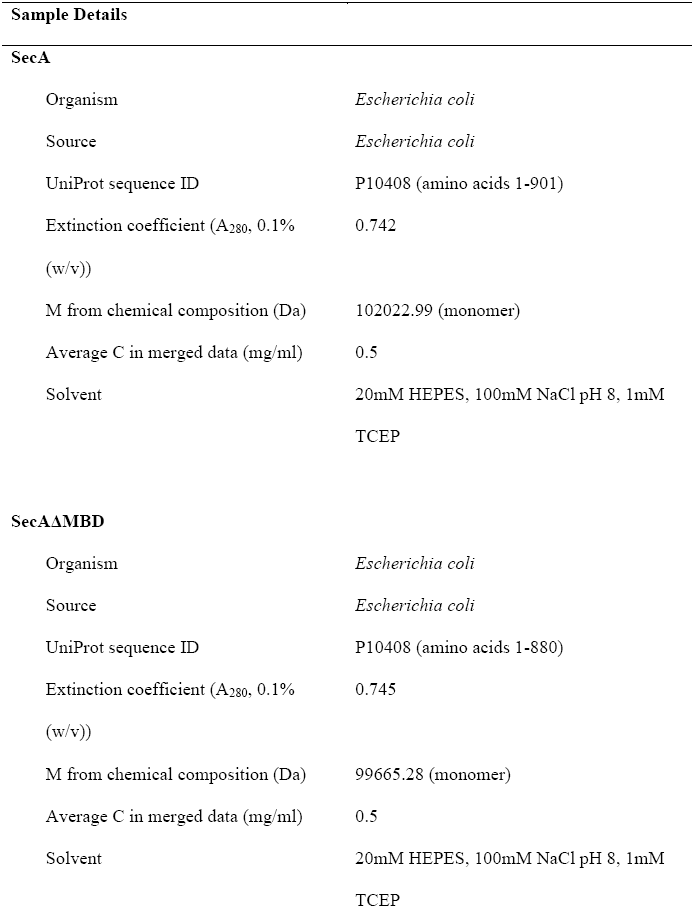
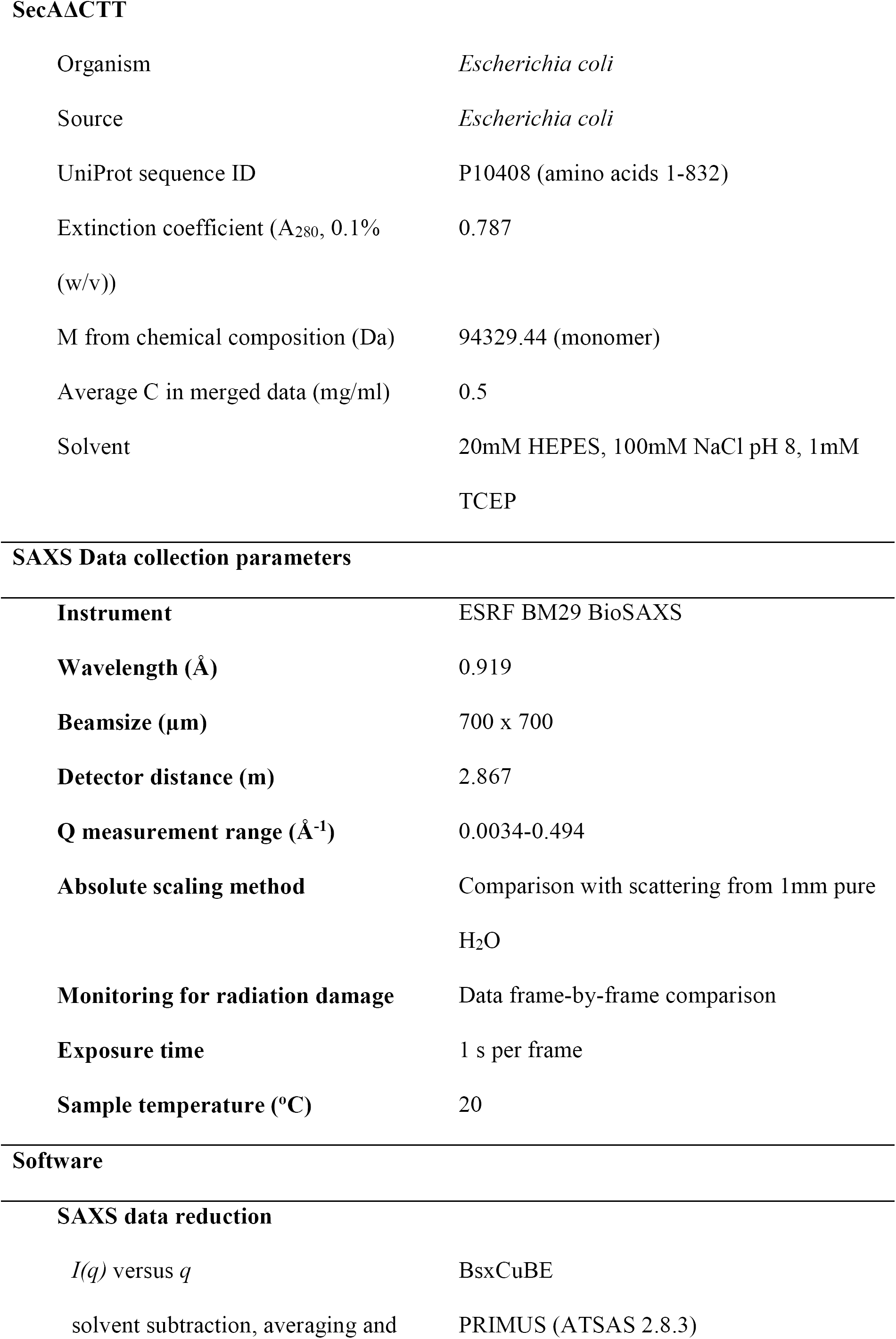
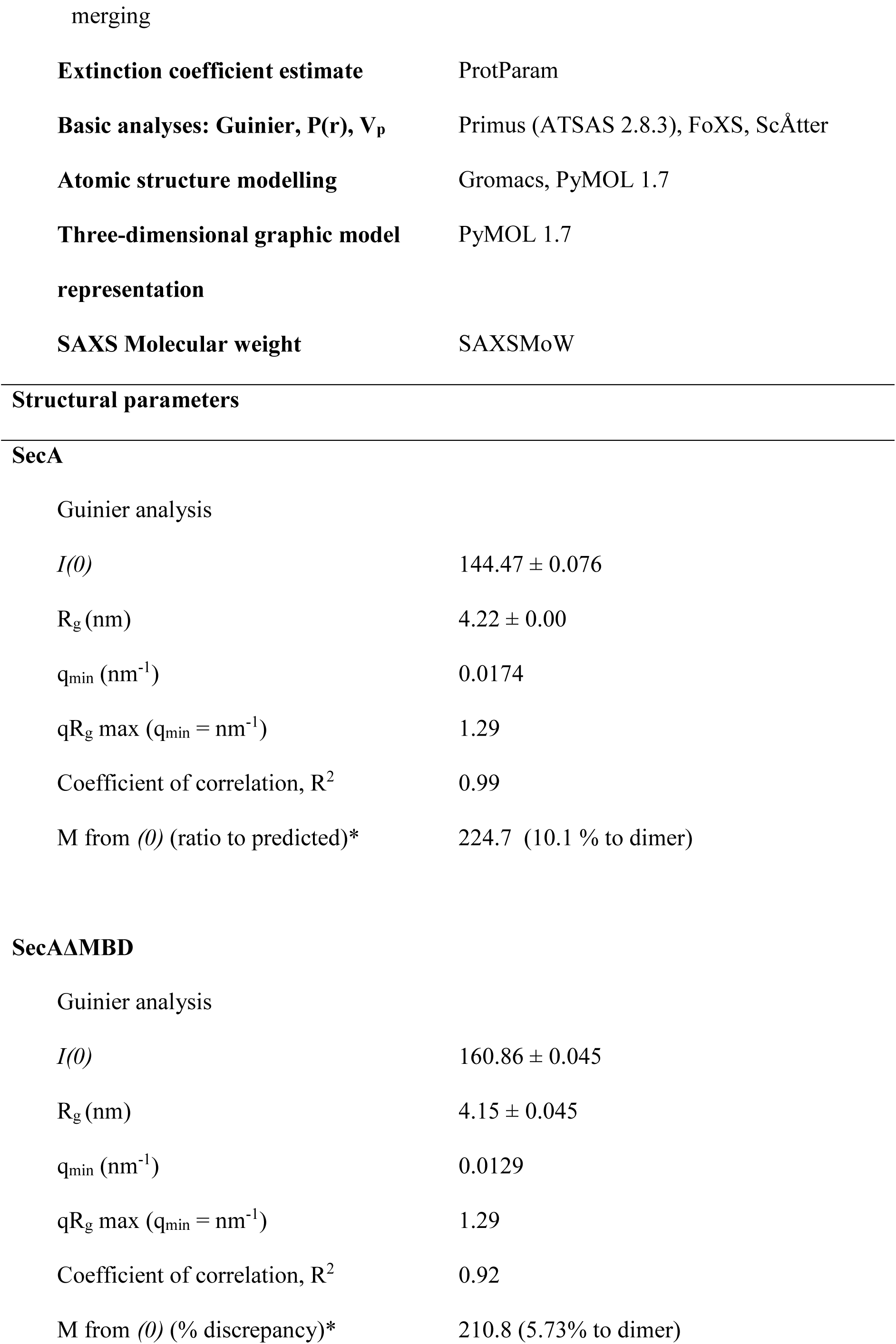
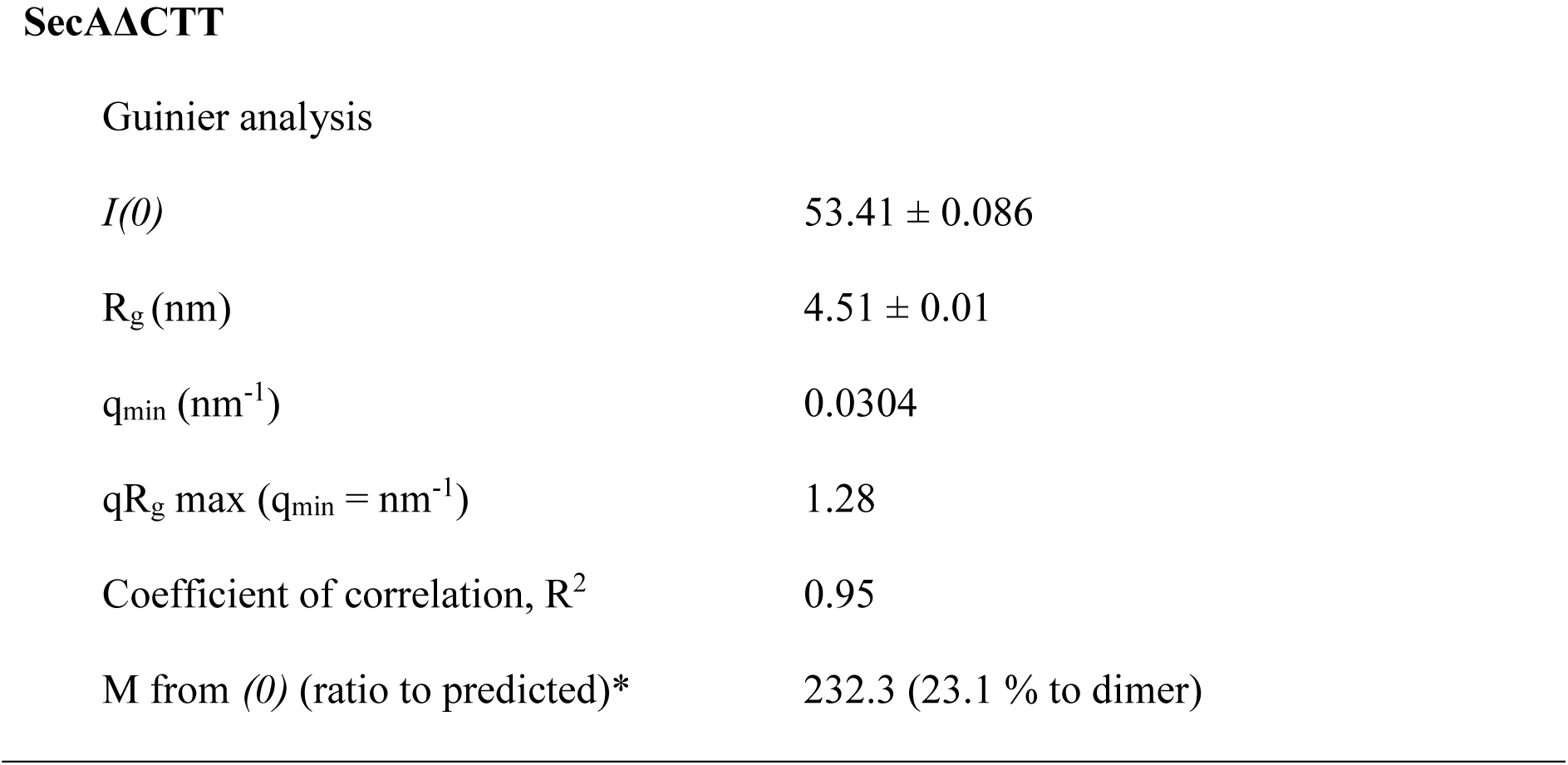
SAXS data collection and processing details for SecA, SecAΔMBD and SecAΔCTT.

|  |  |
| --- | --- |
| Organism | *Escherichia coli* |
| Source | *Escherichia coli* |
| UniProt sequence ID | P10408 (amino acids 1-901) |
| Extinction coefficient ($A_{280}$, 0.1%<br>(w/v)) | 0.742 |
| M from chemical composition (Da) | 102022.99 (monomer) |
| Average C in merged data (mg/ml) | 0.5 |
| Solvent | 20mM HEPES, 100mM NaCl pH 8, 1mM<br>TCEP |
| **SecAΔMBD** |  |
| Organism | *Escherichia coli* |
| Source | *Escherichia coli* |
| UniProt sequence ID | P10408 (amino acids 1-880) |
| Extinction coefficient ($A_{280}$, 0.1%<br>(w/v)) | 0.745 |
| M from chemical composition (Da) | 99665.28 (monomer) |
| Average C in merged data (mg/ml) | 0.5 |
| Solvent | 20mM HEPES, 100mM NaCl pH 8, 1mM<br>TCEP |
| **SecAΔCTT** |  |
| Organism | *Escherichia coli* |
| Source | *Escherichia coli* |
| UniProt sequence ID | P10408 (amino acids 1-832) |
| Extinction coefficient ($A_{280}$, 0.1% (w/v)) | 0.787 |
| M from chemical composition (Da) | 94329.44 (monomer) |
| Average C in merged data (mg/ml) | 0.5 |
| Solvent | 20mM HEPES, 100mM NaCl pH 8, 1mM TCEP |
| **SAXS Data collection parameters** |  |
| **Instrument** | ESRF BM29 BioSAXS |
| **Wavelength (Å)** | 0.919 |
| **Beamsize (μm)** | 700 x 700 |
| **Detector distance (m)** | 2.867 |
| **Q measurement range (Å<sup>-1</sup>)** | 0.0034-0.494 |
| **Absolute scaling method** | Comparison with scattering from 1mm pure H<sub>2</sub>O |
| **Monitoring for radiation damage** | Data frame-by-frame comparison |
| **Exposure time** | 1 s per frame |
| **Sample temperature (°C)** | 20 |
| **Software****SAXS data reduction** |  |
| $I(q)$ versus $q$ | BsxCuBE |
| solvent subtraction, averaging and | PRIMUS (ATSAS 2.8.3) |
| merging |  |
| **Extinction coefficient estimate** | ProtParam |
| **Basic analyses: Guinier, $P(r)$, $V_p$** | Primus (ATSAS 2.8.3), FoXS, ScÅtter |
| **Atomic structure modelling** | Gromacs, PyMOL 1.7 |
| **Three-dimensional graphic model representation** | PyMOL 1.7 |
| **SAXS Molecular weight** | SAXSMoW |
| **Structural parameters** |  |
| **SecA** |  |
| Guinier analysis |  |
| $I(0)$ | $144.47 \pm 0.076$ |
| $R_g$ (nm) | $4.22 \pm 0.00$ |
| $q_{\min}$ (nm<sup>-1</sup>) | 0.0174 |
| $qR_g$ max ($q_{\min} = \text{nm}^{-1}$) | 1.29 |
| Coefficient of correlation, $R^2$ | 0.99 |
| M from $(0)$ (ratio to predicted)* | 224.7 (10.1 % to dimer) |
| **SecAΔMBD** |  |
| Guinier analysis |  |
| $I(0)$ | $160.86 \pm 0.045$ |
| $R_g$ (nm) | $4.15 \pm 0.045$ |
| $q_{\min}$ (nm<sup>-1</sup>) | 0.0129 |
| $qR_g$ max ($q_{\min} = \text{nm}^{-1}$) | 1.29 |
| Coefficient of correlation, $R^2$ | 0.92 |
| M from $(0)$ (% discrepancy)* | 210.8 (5.73% to dimer) |
| $I(0)$ | $53.41 \pm 0.086$ |
| $R_g$ (nm) | $4.51 \pm 0.01$ |
| $q_{\min}$ (nm<sup>-1</sup>) | 0.0304 |
| $qR_g$ max ($q_{\min} = \text{nm}^{-1}$) | 1.28 |
| Coefficient of correlation, $R^2$ | 0.95 |
| M from $(0)$ (ratio to predicted)* | 232.3 (23.1 % to dimer) |

**Supplemental table S3.**
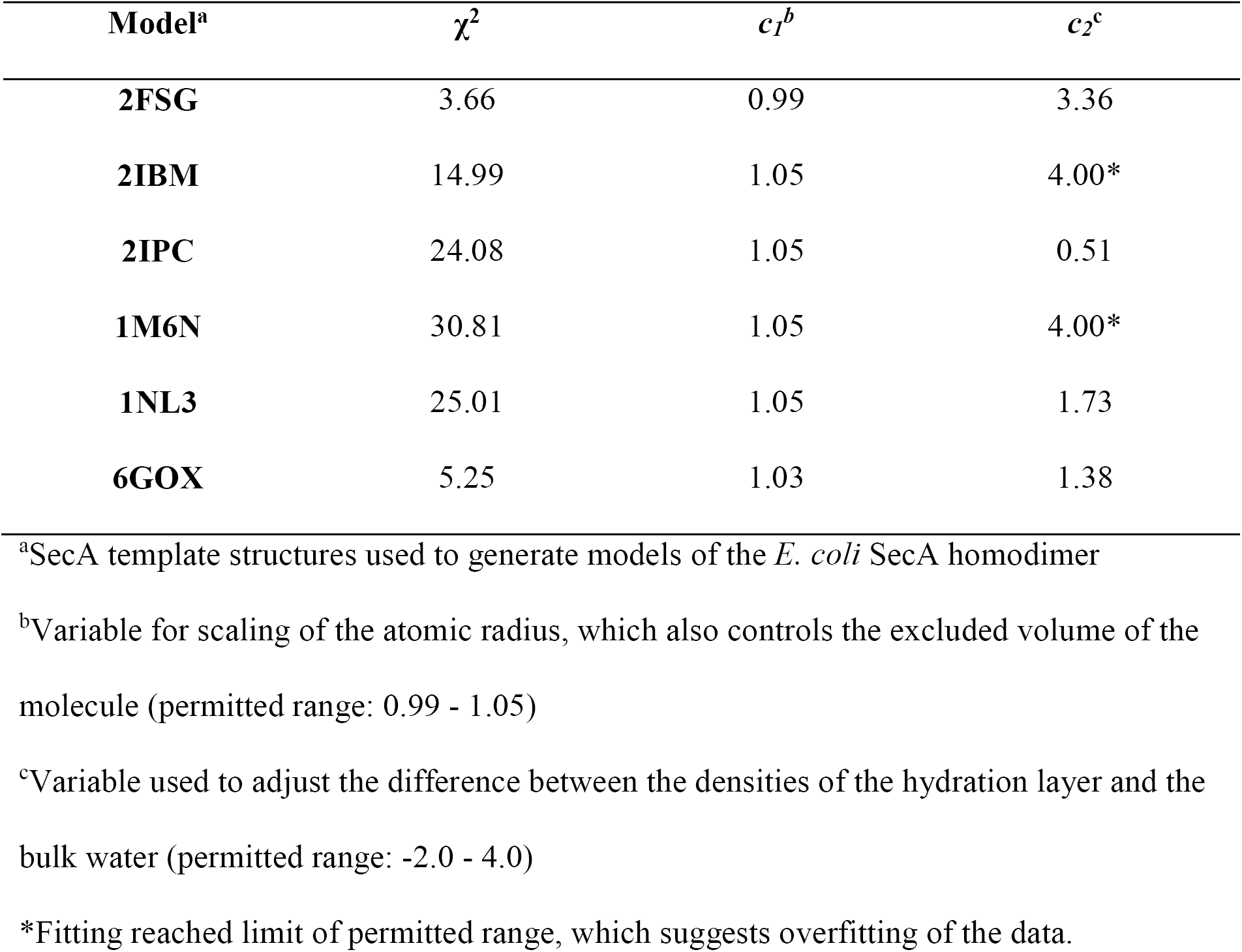
Fitting parameters of models of the *E. coli* SecA dimer.

**Supplemental figure S5 (associated with figure 5).**
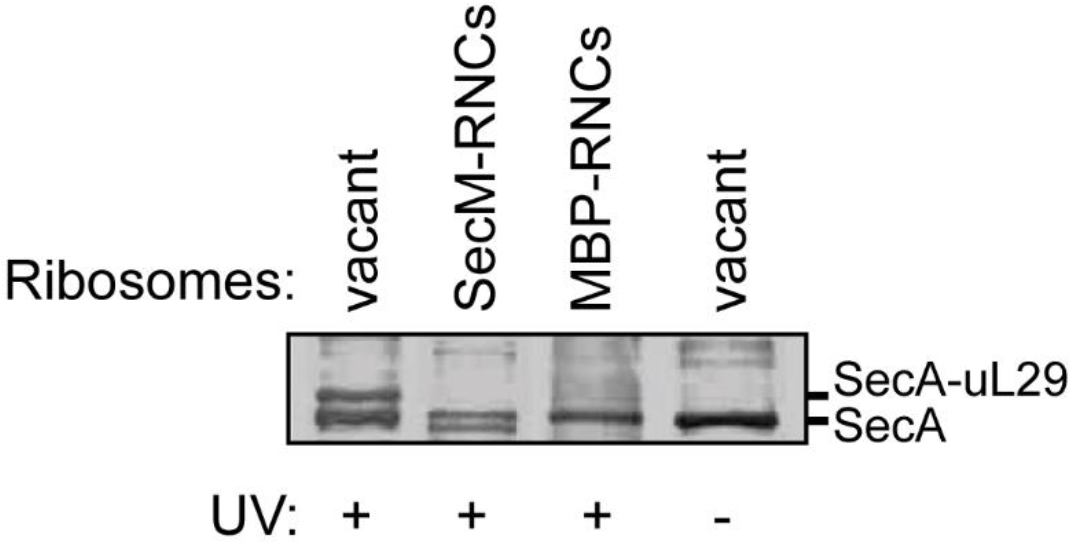
Binding of SecA to the ribosome. 1 μM SecA^Bpa399^ was incubated with non-translating 70S ribosomes (vacant) or RNCs containing arrested nascent SecM (SecM-RNCs) or maltose binding protein (MBP-RNCs). Where indicated, samples were exposed to light at 350 nm (UV). Samples were then resolved using SDS-PAGE and probed by western blotting against SecA. The running positions of SecA and the crosslinking adduct between SecA and ribosomal protein uL29 are indicated.

